# Phylogenomics resolve the systematics and biogeography of the ant tribe Myrmicini and tribal relationships within the hyperdiverse ant subfamily Myrmicinae

**DOI:** 10.1101/2024.08.01.606207

**Authors:** Matthew Prebus, Christian Rabeling

## Abstract

Ants are a globally distributed and highly diverse group of eusocial animals, playing key ecological roles in most of the world’s terrestrial ecosystems. Our understanding of the processes involved in the evolution this diverse family is contingent upon our knowledge of the phylogeny of the ants. While relationships among most subfamilies have come into resolution recently, several of the tribal relationships within the hyperdiverse subfamily Myrmicinae persistently conflict between or within studies, mirroring the controversial relationships of the Leptanillinae and Martialinae to the remaining ant subfamilies. Another persistent issue of debate in ant phylogenetics is the timing of major evolutionary events as inferred via divergence dating. Here, we test the topology of the myrmicine tribes using genome scale data, inspect gene tree-species tree concordance, and use posterior predictive checks and tests of compositional heterogeneity to infer sequence characteristics which potentially introduce systematic bias in myrmicine tribal topology. Furthermore, we test the placement of the fossil †*Manica andrannae* by integrating phylogenomic and morphological data from nearly all species within the genus *Manica,* and a broad sampling of its sister genus *Myrmica.* Subsequently, we demonstrate the effect of fossil placement on overall divergence times in the Myrmicinae. We then re-evaluate the historical biogeography of the Myrmicini and Pogonomyrmecini considering newly generated genetic data and insights from our phylogenomic results. We find that our current understanding of tribal topology in the Myrmicinae is strongly supported, but this topology is highly sensitive to compositional heterogeneity and gene-tree species-tree conflict. Our fossil placement analyses strongly suggest that †*Manica andrannae* is a stem *Manica* species, and that placement of this fossil in the crown group affects not only divergence dates within the tribe Myrmicini, but also has broad implications for divergence times throughout the formicoid clade. The results of our biogeographic reconstructions indicate a South American origin for the Pogonomyrmecini + Myrmicini, with the MRCA of *Myrmica* inhabiting the western Nearctic in the early Miocene prior to repeated dispersal across Beringia throughout the Miocene and Pliocene. The MRCA of *Manica*, on the other hand, was inferred to have a Holarctic range prior to vicariance during the Pliocene. Unexpectedly, we found strong support in the Pogonomyrmecini for three coordinated dispersal events from South to Central America during the early Miocene, which has been previously proposed as an early biotic interchange event prior to the more commonly accepted 3.5 Ma closure of the Isthmus of Panama.

---

Ants are a globally distributed and highly diverse group of eusocial animals, playing key ecological roles in the vast majority of the world’s terrestrial ecosystems (Hölldobler and Wilson, 1990). Roughly half of all extant ant species belong to the subfamily Myrmicinae (7,138 of 14,187 spp.; Bolton 2024), which contains many well-known and ecologically important groups, *e.g.* the leaf-cutter ants (*Atta* and relatives), the fire ants (*Solenopsis*), and the harvester ants (*Messor* and *Pogonomyrmex*). Our understanding of the processes involved in the evolution of this globally successful subfamily is contingent upon our knowledge of their relationships.

With the advent of broad-scale molecular phylogenetic studies, the relationships among most subfamilies have come into resolution (Moreau et al. 2006; Brady et al. 2006; Rabeling et al. 2008; Borowiec et al. 2019; Romiguier et al. 2022; Cai 2024). However, since the six-tribe scheme was proposed by Ward et al. (2015), several of the tribal relationships within the Myrmicinae persistently conflict between or within studies (Ward et al. 2015; Branstetter et al. 2017a; Romiguier et al. 2022). Specifically, the relationships of the tribes Myrmicini, Pogonomyrmecini, Attini, and Solenopsidini remain in question (see Fig. 1) and are either recovered as a grade (Ward el al. 2015; Nelsen et al. 2018; Romiguier et al. 2022: supertree analysis) or as two clades composed of [Myrmicini + Pogonomyrmecini] and [Attini + Solenopsidini] (Branstetter et al. 2017a; Romiguier et al. 2022: supermatrix analysis). Intriguingly, the pattern of recovering a grade vs. clade for Myrmicini + Pogonomyrmecini, both of which are subtended by long branches, mirrors the conflicting topologies recovered for Leptanillinae + Martialinae (Rabeling et al. 2008; Borowiec et al. 2019; Romiguier et al. 2022; Cai 2024).

**Figure 1.**
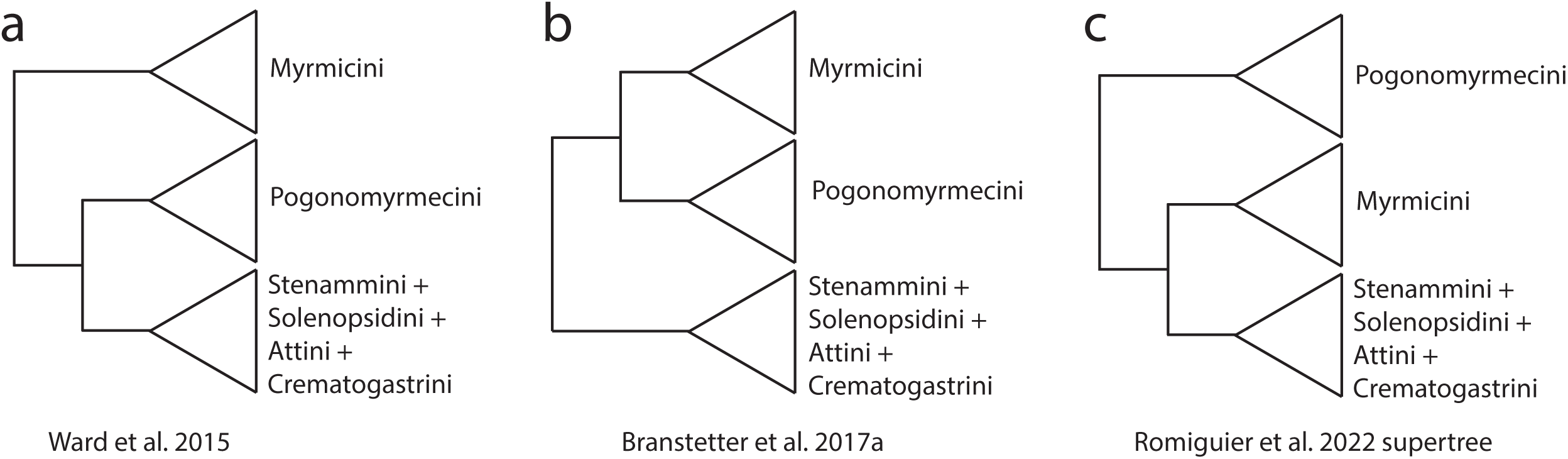
Tribal topologies of the ant subfamily Myrmicinae inferred in previous studies.

Another persistent issue of debate in ant phylogenetics is the timing of major evolutionary events as inferred via divergence dating analyses (Borowiec et al. 2021). Recent studies have shown that many fossils previously assigned to specific taxa have been either erroneously placed, or their position becomes ambiguous upon scrutiny (Boudinot et al. 2022a & b). Following a revelation about the species status of *Manica parasitica* (Prebus et al. 2023) and inspired by recent discoveries of Eocene *Manica* fossils (Zharkov et al. 2022; LaPolla 2023), the focal point of this study is the systematics and historical biogeography of the ant genus *Manica* and its parent tribe the Myrmicini. We test the impact of fossil placement in *Manica,* because one fossil which has been proposed as a member of the crown group (i.e. within the clade of extant species) influences how the timing of evolutionary events in the subfamily Myrmicinae is interpreted (Zharkov et al. 2022).

Previous biogeographical analyses of *Myrmica*, the speciose sister clade of *Manica*, have been confounded by poor phylogenetic resolution (Jansen et al. 2010). While there have been multiple hypotheses extended for the geographical origin and biogeographical history of *Myrmica, Manica,* and the Myrmicini (Wheeler & Wheeler 1970; Radchenko & Elmes 2001; Jansen & Savolainen 2010; Zharkov et al. 2022), to date these aspects of their evolutionary history remain murky, either with ambiguous results (Jansen et al. 2010) or with sparse species sampling (Ward et al. 2015).

Here, we test the placement of the fossil †*M. andrannae* proposed by Zharkov et al. (2022) by integrating phylogenomic and morphological data from nearly all species within the genus *Manica,* and a broad sampling of its sister genus *Myrmica.* While †*M. andrannae* bears the diagnostic characters that unite extant *Manica* species, the well-developed propodeal tubercles, narrow head capsule, and short propodeum observed in †*M. andrannae* are divergent from all extant *Manica*, and closer to the conditions observed in extant *Myrmica.* Subsequently, we demonstrate the effect of fossil placement on overall divergence times in the Myrmicinae. Furthermore, we test the topology of the myrmicine tribes using genome scale data, inspect gene tree-species tree concordance, and use posterior predictive checks and tests of compositional heterogeneity to infer sequence characteristics which potentially introduce systematic bias in the tribal topology. Finally, we re-evaluate the historical biogeography of the Myrmicini and Pogonomyrmecini considering newly generated genetic data and insights from our phylogenomic results.

## Materials & Methods

### Material Examined

Because †*Manica iviei* LaPolla is a compression fossil of an alate gyne, many morphological characters are not visible. Additionally, we had very few examples of extant gynes available to us; therefore, we excluded †*M. iviei* from this study. To test the placement of the fossil †*Manica andrannae* Zharkov et al. within the tribe Myrmicini, we gathered specimens of all extant *Manica* species, including samples from across the geographic range of each species whenever possible. Additionally, we included multiple species from the genus *Myrmica*, which constitutes the other genus in the Myrmicini, including as many lineages as possible, basing our sampling on Jansen et al. (2010); we included multiple species from the genus *Pogonomyrmex* as well. In total, our dataset includes 59 new samples. Collection data for new samples used in this study can be found in Supplementary Table 1. Voucher specimens were deposited in the repositories designated in the “OwnedBy” field in Supplementary Table 1, using the following repository codes:

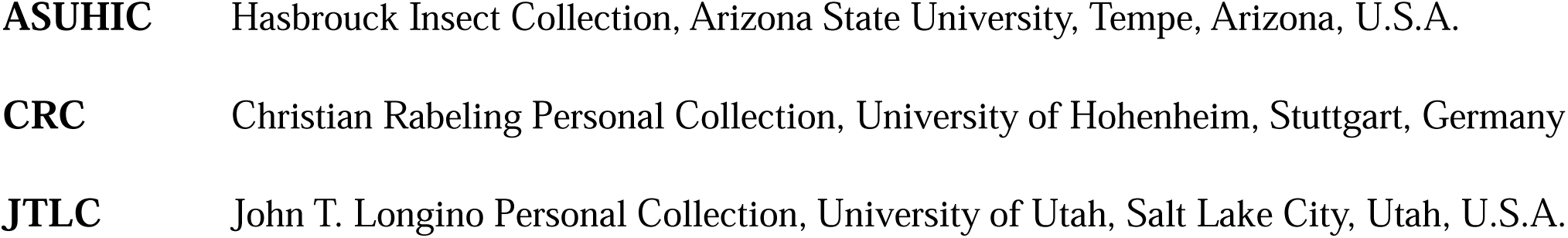
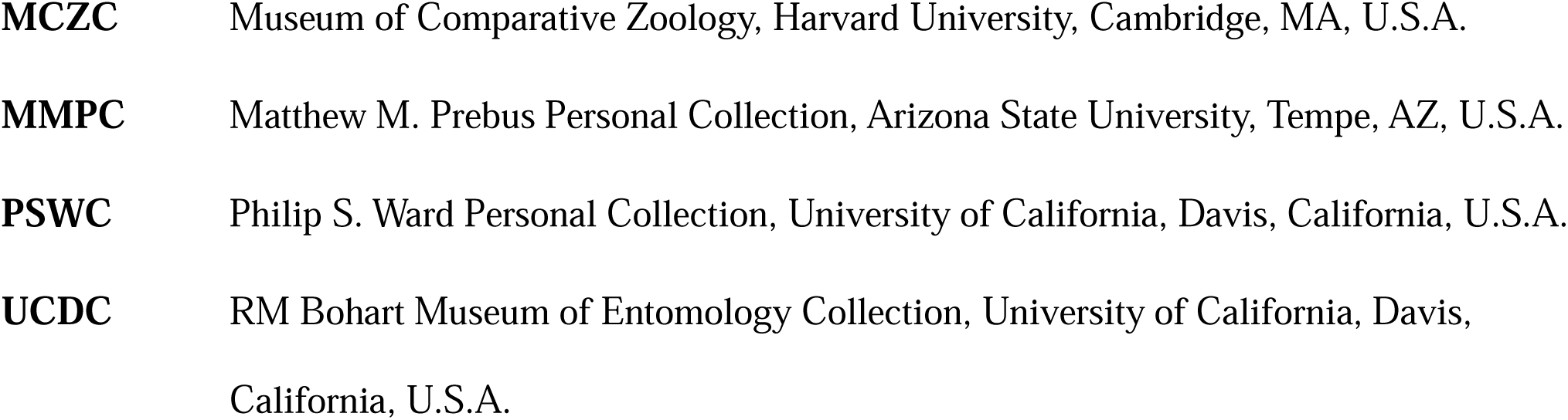

### Molecular Phylogeny

#### Molecular data generation

To generate the genetic data, we extracted DNA, prepared genomic libraries, and performed targeted enrichment of ultraconserved elements (UCEs) using a set of RNA probes targeting 2,524 UCE loci in the Hymenoptera (Branstetter et al. 2017a). We extracted DNA nondestructively from adult worker ants by using a flame-sterilized size 2 stainless steel insect pin to pierce the cuticle of the head, mesosoma, and gaster on the right side of the specimens, then used a DNeasy Blood and Tissue Kit (Qiagen, Inc., Hilden, Germany) following the manufacturer’s protocols. We verified DNA extract concentration using a Qubit 3.0 Fluorometer (Invitrogen, Waltham, MA, U.S.A.). We input up to 50 ng of DNA, sheared to a target fragment size of 400–600 bp with a QSonica Q800R sonicator (Qsonica, Newtown, CT, U.S.A.) into a genomic DNA library preparation protocol (KAPA HyperPrep Kit, KAPA Biosystems, Wilmington, MA, U.S.A.) for targeted enrichment of UCEs following Faircloth et al. (2015) as modified by Branstetter et al. (2017a) using a unique combination of iTru barcoding adapters (Glenn et al. 2019; BadDNA, Athens, GA, U.S.A.) for each sample. We performed enrichments on pooled libraries using the catalog version of the Hym 2.5Kv2A ant-specific RNA probes (Branstetter et al., 2017a; Arbor Biosciences, Ann Arbor, MI, U.S.A.), which targets 2524 UCE loci in the Formicidae. We followed the library enrichment procedures for the probe kit, used custom adapter blockers instead of the standard blockers (Glenn et al. 2019; BadDNA, Athens, GA, U.S.A.), and left enriched DNA bound to the streptavidin beads during PCR, as described in Faircloth et al. (2015). Following post-enrichment PCR, we purified the resulting pools using SpeedBead magnetic carboxylate beads (Rohland & Reich 2012; Sigma-Aldrich, St. Louis, MO, U.S.A.) and adjusted their volume to 22 μl. We verified enrichment success and measured size-adjusted DNA concentrations of each pool with qPCR using a SYBR-FASTqPCR kit (Kapa Biosystems, Wilmington, MA, U.S.A.) and a Bio-Rad CFX96 RT-PCR thermal cycler (Bio-Rad Laboratories, Hercules, CA, U.S.A.) and combined all pools into an equimolar final pool. We sequenced the final pool at Novogene (Sacramento, CA, U.S.A.) on Illumina HiSeq 150 cycle Paired-End Sequencing v4 runs (Illumina, San Diego, CA, U.S.A.), along with other enriched libraries for unrelated projects. Raw reads can be found on NCBI (BioProject PRJNA1072056).

#### Molecular data processing

We processed the resulting raw reads with the PHYLUCE pipeline (Faircloth 2016), by first removing adapter contamination and low-quality reads using *illumiprocessor,* then assembling the cleaned reads with SPAdes v3.15.3 (Prjibelski et al. 2020).

#### Dataset construction: Myrmicini UCEs

To increase our ingroup and outgroup sampling, we incorporated sequences from previously published datasets (Blaimer et al. 2018; Branstetter et al 2017a; Branstetter et al 2017b; Prebus et al. 2023; Doering et al. 2024), yielding a final dataset of 62 samples.

We followed the standard PHYLUCE protocol for processing UCEs in preparation for phylogenomic analysis, aligning the monolithic unaligned FASTA file with the phyluce_align_seqcap_align command, using MAFFT (Katoh & Standley 2013) as the aligner (—aligner mafft) and opting not to edge-trim the alignment (–no-trim). We trimmed the resulting alignments with the phyluce_align_get_gblocks_trimmed_alignments_from_untrimmed command in PHYLUCE, which uses GBlocks ver. 0.91b (Castresana 2000), using the following settings: b1 0.5, b2 0.5, b3 12, b4 7. After removing UCE locus information from taxon labels using the command phyluce_align_remove_locus_name_from_nexus_lines, we examined the alignment statistics using the command phyluce_align_get_align_summary_data and generated a dataset in which each locus contains a minimum of 85% of all taxa using the command phyluce_align_get_only_loci_with_min_taxa.

#### Phylogenetic inference

After an initial phylogeny construction with IQTREE v2.1.1 (Minh et al. 2020) and inspection of the alignment with AliView (Larsson 2014), we found that branch lengths for some taxa were longer than expected due to poor data quality. To correct this, we used the SPRUCEUP pipeline (Borowiec 2019), with an overall lognormal cutoff of 0.97, and manual cutoffs as follows:

a. Manica_hunteri_CA_Coffee_Creek_P590: 0.09;
b. Manica_hunteri_WA_Bean_Creek_P591: 0.09;
c. Manica_rubida_CZ_P602: 0.08;
d. Manica_invidia_OR_Alkali_Lake_P594: 0.09.

Because the assumption that the evolutionary rates of sequence data are homogenous is often violated in empirical data (Buckley et al. 2001), we partitioned our UCE loci into sets of similarly evolving sites. To achieve this, we used the command phyluce_align_format_nexus_files_for_raxml which concatenates loci into a single alignment and generates a partition file for input into the SWSC-EN method (Tagliacollo & Lanfear 2018). We used the resulting datablocks as input for partitioning in IQTREE2, using the command -m MFP + MERGE. Because the combination of gamma and proportion of invariable sites (+I+G) has been demonstrated to result in anomalies in likelihood estimation (Sullivan & Swofford 2001; Yang 2006), we set the rate heterogeneity models to a subset that includes everything except the combination of gamma and proportion of invariable sites (-mrate E, I, G). We set the search algorithm to -rclusterf 10. We used the resulting partitioned dataset as input for maximum likelihood tree inference in IQTREE2 using 1000 ultrafast bootstrap replicates (-bb 1000).

Additionally, we performed a summary coalescent species tree analysis with ASTRAL III v5.7.4 (Zhang et al. 2018). As input, we used individual locus trees calculated using the same partition scheme and settings in IQTREE2, as above.

### Morphology

#### Protocol for morphometric data collection

To evaluate the placement of †*Manica andrannae* in the Myrmicini, we gathered morphometric data from all *Manica* and *Myrmica* molecular voucher specimens in our dataset using the measurements collected by Zharkov et al. (2022) as a guideline. To take our measurements, we used a Leica M205C stereo microscope and a moveable stage equipped with orthogonal digital micrometers. Morphometric data have been deposited in MorphoBank (http://morphobank.org/permalink/?P5425). The following measurements were taken (see Supplementary file 1, Fig. S1 for an illustration of measurements):

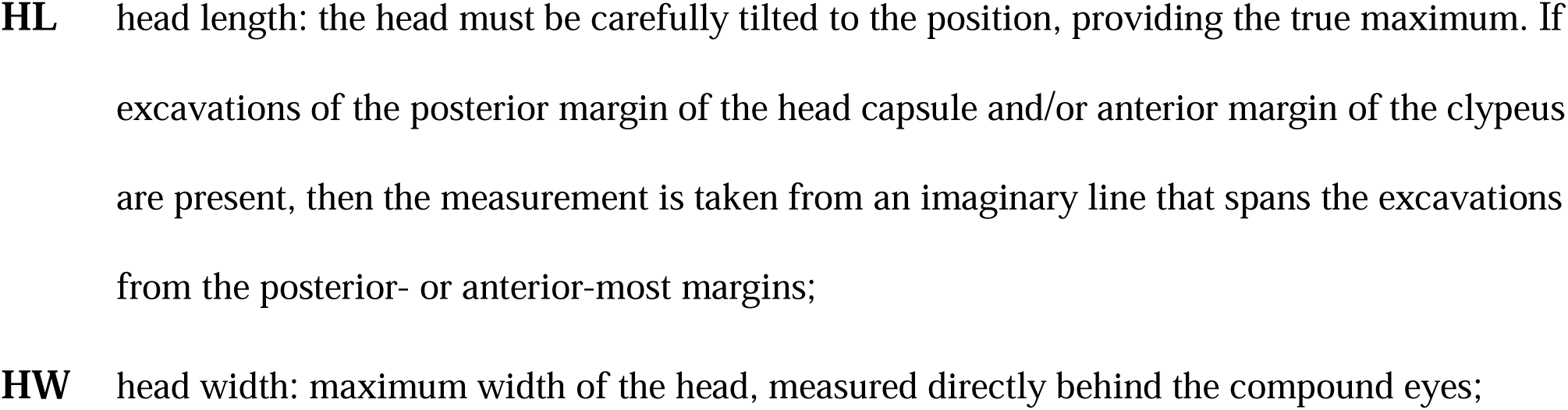
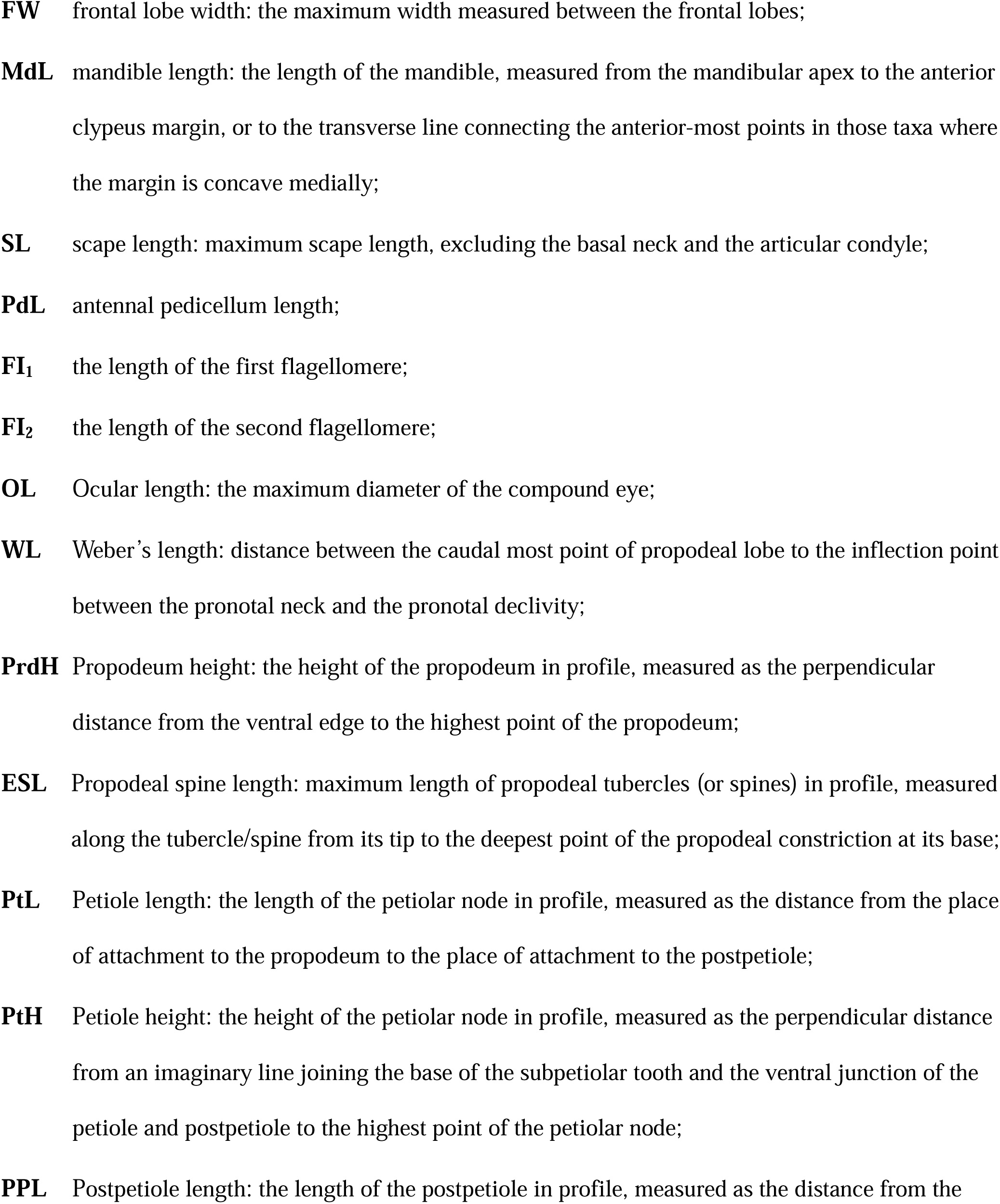
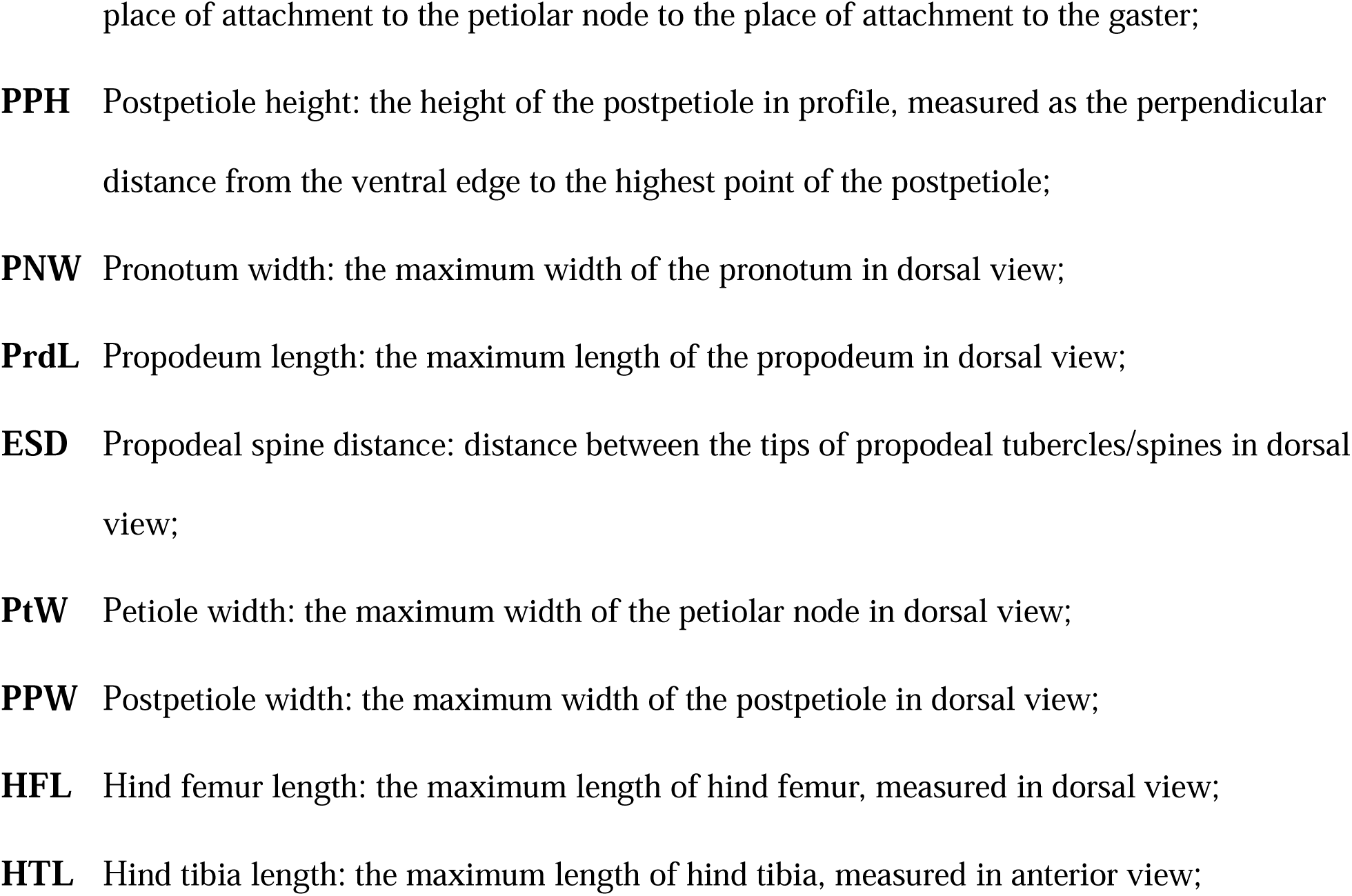

The following indices were used:

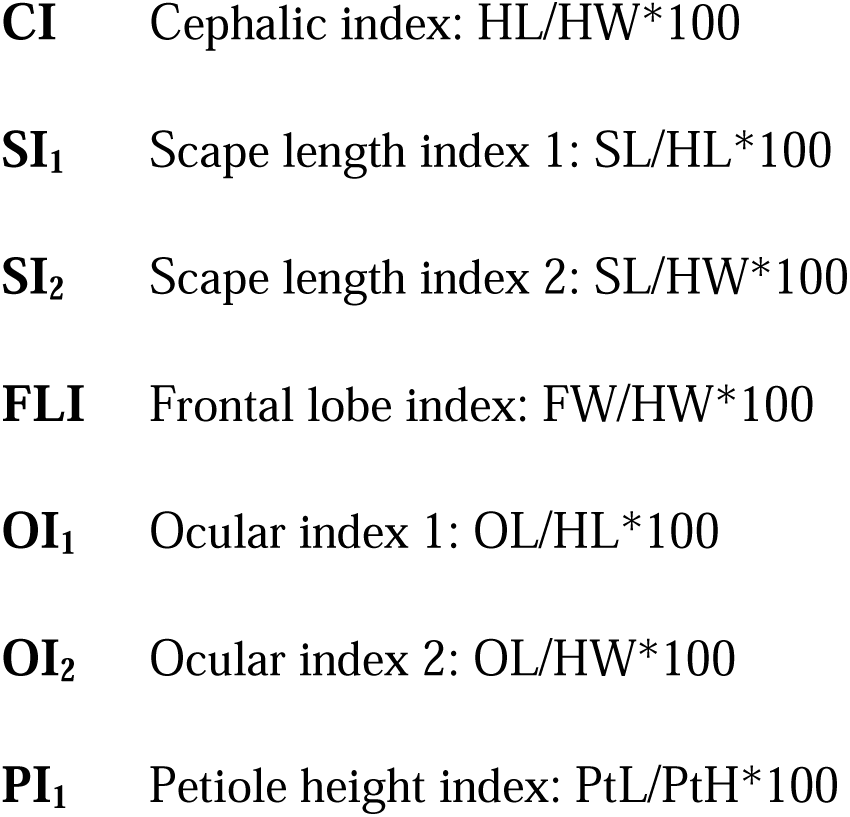
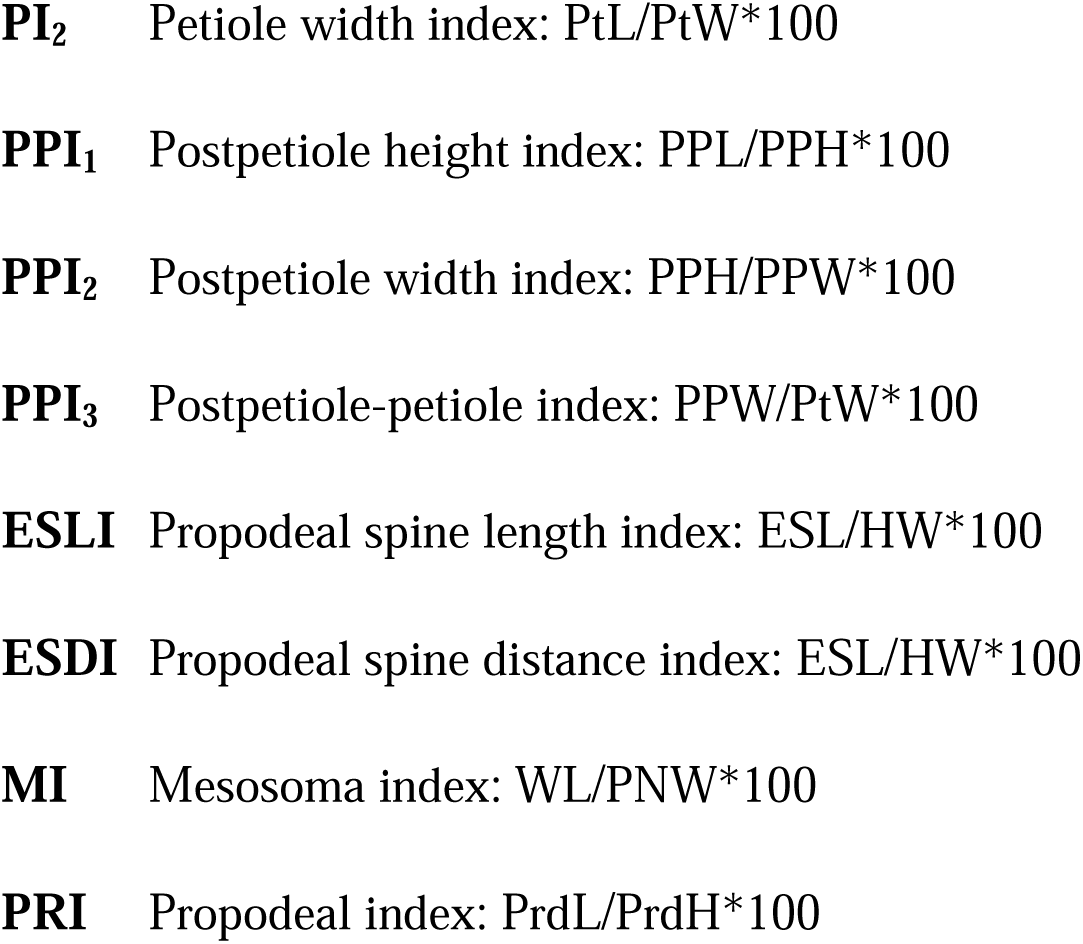

#### Analysis of morphometric data

For an initial inspection of our data, we used ggplot2 in R v4.1.3 (Wickham 2016; R Core Team 2022) to plot and compare the overall distributions of each measurement and index taken from extant *Manica* and *Myrmica* against the measurements of †*Manica andrannae* derived from Zharkov et al. (2022). We compared the averages of measurements of *Man ca* against *Myrmica* using a *t-test* at a significance level of *p* < 0.05. To translate the data into binary character states in preparation for the fossil placement analysis, we first calculated the medians of each measurement or index for extant *Manica* and extant *Myrmica.* Then, we calculated the distance between the medians as:

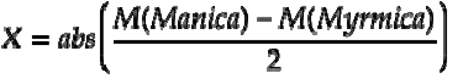

We then found the midpoint of the distance by adding the distance to the smaller of M(*Manica*) vs. M(*Myrmica*). Following this, measurements for each voucher were coded with the following logic: if a measurement for a given voucher was greater than the midpoint, it was coded as ‘1’; otherwise, it was coded as ‘0’.

### *Manica* Fossil Placement and Divergence Dating

#### Dataset construction: reduced Myrmicini UCEs

To accommodate computational limitations of the fossil placement analysis, we used SortaDate (Smith et al. 2018) with the individual locus trees inferred above as input to reduce the full dataset to a set of 50 UCE loci, sorting the trees first by bipartition, then by root-to-tip variance, then by tree length.

#### Phylogenetic inference and divergence dating

To compare our reduced dataset against the full dataset, we inferred a phylogeny using IQTREE2 using the same settings as the previous analysis, but used the ‘mrbayes’ model set (-mset mrbayes) in order to restrict the substitution model search to those used by MRBAYES only. We then designed three analyses for MRBAYES v3.2.6 (Ronquist et al. 2012):

1. Morphology only: we evaluated the placement of †*Manica andrannae* based on morphology alone in a non-clock analysis, using two runs with four chains each, sampling every 2500 generations for 150 M generations.
2. Molecules only: to evaluate the topology of the phylogeny inferred from our molecular dataset, we used the partitioning scheme found by the IQTREE2 analysis above in a non-clock analysis. We then analyzed the dataset with two runs with four chains each, sampling every 2500 generations for 5 M generations.
3. Combined morphology and molecules: to evaluate the topology of the phylogeny inferred from the combined dataset, we used the IQTREE2 partitioning scheme in a non-clock analysis, analyzing the dataset with two runs with four chains each, sampling every 2500 generations for 5 M generations.

We then performed three alternate time calibrated analyses on the combined dataset: an unconstrained analysis, and two constrained analyses comparing the topology suggested by Zharkov et al. (2022) and the topology suggested by the morphology-only and unconstrained analyses above. For each of these last three analyses, we used the fossilized birth-death branch lengths prior (brlenspr = clock:fossilization), and the IGR clock variation prior (clockvarpr = igr). We used a uniform calibration on the root node, with a range of 45.3–61 Ma based on Ward et al. (2015) (treeagepr = uniform(45.3,61)), and a uniform calibration with a range of 33.9–37.8 Ma for †*Manica andrannae* based on Perkovsky et al. (2007) (uniform(33.9,37.8)). We set the sampling probability to 0.1 based on the number species in our dataset vs. the number of extant Myrmicini (18/192 spp). For each of these analyses, we used two runs with four chains each, sampling every 2500 generations for 150 M generations.

### *Manica* Fossil Placement Effects on Myrmicinae Divergence Dating

To investigate the downstream effects of alternate fossil placements in the Myrmicini on the overall divergence times within Myrmicinae, we performed divergence dating analyses on a broader sampling of myrmicines and outgroups.

#### Dataset construction: Myrmicinae UCEs

We reduced the full Myrmicini dataset by removing replicate population samples of *Manica* species, using the number of UCE loci recovered as a criterion.

We then incorporated a broader sampling of the Myrmicinae and other formicoid clade outgroups from previous studies (Prebus 2017; Branstetter et al. 2017a; Branstetter et al. 2017b; Pierce et al. 2017; Blaimer et al. 2018; Schifani et al. 2022). We followed the protocol described above for the ‘Myrmicini UCEs’ dataset construction.

#### Phylogenetic inference and divergence dating

We partitioned our data, then inferred maximum likelihood and summary coalescent phylogenies as above in the ‘Myrmicini UCEs’ analyses. To reduce the computation time for the divergence dating analysis in MCMCTREE (Yang 1997, 2007), we reduced the full Myrmicinae UCE dataset by using the gene trees from the summary coalescent analysis as input for SortaDate, selecting 50 loci based on the sorting criteria used in the Myrmicini divergence dating analysis. In a further effort to reduced computation time, we used the approximate likelihood calculation (Reis & Yang 2011). We used the summary coalescent topology from the ASTRAL analysis above and compared divergence estimates using 31 fossil calibrations (see Supplementary Table 2) and two alternate calibrations in the Myrmicini:

1. Crown *Manica* calibrated with > 44.1 Ma (this assumes that †*Manica andrannae* is part of the *Manica* crown group; we used the age of Baltic amber from Constenius 1996), and a calibration of >44.1 Ma for the Myrmicini based on Baltic amber *Myrmica* fossils.
2. Only Myrmicini calibrated with > 44.1 Ma (this assumes that †*Manica andrannae* is part of the *Manica* stem group).

We used a secondary calibration for the root, < 120 Ma based on the age of the formicoid clade from Borowiec et al. (2021). We set the clock to the independent rates model (clock = 2) and used the HKY85 substitution model (model = 4). We set the birth-death prior (BDparas) to 1 1 0.1. We set the gamma prior for overall substitution rate for genes (rgene gamma) to 1 5.7 using the results from an initial clock analysis in baseml. Finally, we set the number of samples to 10,000 with a sample frequency of 5,000 and a burnin of 5,000,000. We ran two analyses in parallel to confirm that results were stable and checked for convergence using Tracer v1.7.2 (Rambaut et al. 2018). We summarized the results of the analysis using the R package MCMCtreeR (Puttick 2019).

### Tribal Topology

*Topology tests.—* To test the tribal topology of the Myrmicinae, specifically focusing on the placement of the Myrmicini, we used the full sequence dataset and from above and constructed alternate constrained tree topologies to use as input into IQTREE2:

1. Ward et al. 2015 & Nelsen et al. 2018 (Fig. 1a)
2. Branstetter et al. 2017a, Blaimer et al. 2018, and Romiguier et al. 2022 concatenate and partition (supermatrix) analysis (Fig. 1b)
3. Romiguier et al. 2022 summary coalescence (supertree) analysis (Fig. 1c)

We compared these trees using five tree topology tests: RELL approximation (Kishino et al. 1990); the Kishino-Hasegawa test (Kishino and Hasegawa, 1989); the Shimodaira-Hasegawa test (Shimodaira and Hasegawa, 1999); expected likelihood weights (Strimmer and Rambaut, 2002); and the approximately unbiased test (Shimodaira, 2002). We set the number of RELL replicates to 10,000.

#### Gene tree/species tree discordance

To investigate gene tree/species tree discordance as a factor in topological conflict between analyses, we used gene trees estimated from the full dataset as input into gene concordance factor analysis in IQTREE2, testing them against each of the three topologies above.

#### Model adequacy

To further investigate which dataset characteristics may potentially influence tree topology, we used ModelFinder in IQTREE2 against each dataset partition found by the SWSC-EN pipeline, limiting the nucleotide substitution models to a set used by RevBayes v1.2.1 (Höhna et al. 2014 & 2016) by using the commands ‘-mset F81,GTR,HKY,JC,K80,K81,TN,TIM,TVM’ and ‘-mrate I,G’. We preformed posterior predictive checks against each partition using the model selected by ModelFinder with a set of custom RevBayes scripts, adapted from Höhna et al. (2018).

Using the midpoint test results with thresholds of p = 0.025 and p = 0.975, we calculated significance for each test statistic for each partition. For each of the three topologies above, we used the results of the concordance factor analysis and found each concordant locus that had at least one partition that was adequately modeled. For each alternate topology, we then found the ratio of these loci to loci that contained no adequately modeled partitions. We made the same calculation for discordant loci and subtracted each (the ratio of adequately modeled loci in concordant trees vs. the ratio of adequately modeled loci in discordant trees) from the overall dataset ratio to visualize which test statistics had the most disparity between concordant and discordant trees for each topology. For each topology we selected the test statistic with the greatest disparity, and constructed datasets from partitions that were adequately modeled vs. inadequately modeled. For each dataset constructed above, we then inferred trees using:

1. the unpartitioned matrix;
2. the partitioned matrix with the ModelFinder models for each partition;
3. a summary coalescent tree using the individual partition trees.

Furthermore, we tested each partition with p4 (Foster 2004) to specifically inspect whether our data significantly violated the assumption of compositional homogeneity across taxa. For each partition, we employed two tests:

1. a chi-squared test for compositional heterogeneity;
2. a phylogeny corrected simulation-based test.

For the latter test, we constructed a BioNJ tree (Gascuel 1997), onto which we simulated 1000 sequence data replicates using the GTR + 4Γ substitution model. Then, the empirical sequence data composition distributions were compared against the simulated data distributions. For each of the datasets, we inferred trees from:

1. an unpartitioned supermatrix;
2. a supermatrix partitioned by SWSC-EN data block;
3. a supertree without collapsing low bootstrap nodes in SWSC-EN trees;
4. a supertree in which nodes in SWSC-EN trees with less than 10 bootstrap support were collapsed into polytomies.

Additionally, we computed sequence frequency statistics for each taxon using AMAS (Borowiec 2016) and mapped them onto the results of the partitioned supermatrix analysis of the full dataset using the ‘fastAnc’ function in the R package phytools v2.1-1 (Revell 2012).

### Biogeography

#### Dataset construction: Myrmicini & Pogonomyrmecini legacy loci

We compiled a dataset of six loci from previously published data (Ward & Downie 2005; Brady et al. 2006; Schultz & Brady 2008; Jansen & Savolainen 2010; Jansen et al. 2010; Demarco & Cognato 2015; Ward et al. 2015; Johnson & Moreau 2016; Supplementary Table 3) and extracted legacy loci from the Myrmicini & Pogonomyrmecini contigs assembled in the above section ‘Molecular Phylogeny’ by using the PHYLUCE commands ‘phyluce_assembly_match_contigs_to_probes’ using a legacy-locus-only bait set and ‘phyluce_assembly_match_contigs_to_barcodes’ to extract the barcoding region of COI. We partitioned each locus by exon/intron, and further partitioned each exon by codon position. We used these partitions as input into IQTREE2 for substitution model selection and gathered similarly evolving partitions together with the commands ‘-m MFP+MERGE’. We limited the substitution models to a subset used by RevBayes v1.2.1 by using the commands ‘-mset F81,GTR,HKY,JC,K80,K81,TN,TIM,TVM’ and ‘-mrate I,G’.

### Tree model

We used the partitioning scheme selected by IQTREE2 as input for RevBayes. For the following, we adapted and integrated several tutorials from Höhna (2021). For the divergence dating sub-model, we used a birth-death tree prior, setting the sampling probability *rho* to 181/318 = 0.58 (species in dataset)/(described species in Myrmicini + Pogonomyrmecini). Because the results of the IQTREE analysis and an unconstrained RevBayes analysis conflicted with the topology of our phylogenomic results (see Supplementary file 2, Fig. S16 & S17) and because we use the crown age of the Myrmicini as a secondary calibration, we set several constraints:

1. Pogonomyrmecini, as an outgroup to Myrmicini;
2. *Patagonomyrmex* + *Hylomyrma*, to make the topology consistent with our phylogenomic results;
3. Myrmicini, to set a calibration for the crown Myrmicini;
4. *Manica hunteri* + *Manica rubida*, to make the topology consistent with our phylogenomic results.

We set the root age to a uniform distribution covering 81.8–113.5 Ma, based on the 95% CI from the ‘Myrmicini fossil placement effects on Myrmicinae divergence dating’ analysis above. Similarly, we set the age of the Myrmicini to a uniform distribution covering 41.6–60.9 Ma based on the 95% CI from the ‘Myrmicini fossil placement and divergence dating’ analysis above. Additionally, we set the molecular clock to an uncorrelated lognormal clock.

### Biogeography model

We assembled a biogeography model which incorporates a time-stratified epoch analysis, considering tectonic movements relative to current biogeographic realms over the evolutionary history of the Myrmicini + Pogonomyrmecini. We discretized the ranges of Pogonomyrmecini + Myrmicini into eight areas: South America, Central America, Caribbean, Western Nearctic, Eastern Nearctic, Western Palearctic, Eastern Palearctic, and Indomalayan (see Fig. 4 inset). Because contemporary species are only known to inhabit a maximum of two biogeographic areas, we limited the maximum number of areas that a species could inhabit to two. To make the epoch sub- model, we split the timeline into ten epochs. We created connectivity matrices for each epoch based upon Scotese (2010), taking into account the timing land bridge formation between South and Central America, and the Beringian and Thulean land bridges between the Nearctic and Palearctic. We ran two alternative analyses, with and without the potential for overwater dispersal. For the overwater dispersal analysis, we created new connectivity matrices and added distance matrices for each epoch, again based upon Scotese (2010). To choose between these alternate models, we ran stepping stone analyses and calculated Bayes factors (Rass & Kafferty 1995) from marginal likelihood estimates.

## Results & Discussion

### Molecular phylogeny: Myrmicini UCEs

To build a backbone phylogeny for testing fossil placement and testing topological hypotheses within the Myrmicini, we generated a phylogenomic dataset of UCEs from representatives of all extant *Manica* species and targeted sampling of *Myrmica*. Following sequencing, assembly, and *in silico* extraction of UCE loci, we recovered an average coverage depth of 63.6x (range: 19.3–104.4x) and a mean contig length of 1,350 bp. For additional assembly statistics, see Supplementary Table 4. After alignment, trimming, and filtering, the resulting matrix was 2.8 Mb long and contained 2,177 UCE loci, with 21.5% gaps or missing data.

To infer the overall topology and statistical support for our backbone phylogeny, we conducted two types of analyses: maximum likelihood (ML) analysis on concatenated and partitioned alignment, and summary coalescent (SC) analysis on a dataset of phylogenies inferred from analysis of single UCE loci. The ML phylogeny was statistically well supported, with all species and species relationships within *Manica* receiving full bootstrap support. Similarly, most of the species relationships within *Myrmica* received full support, with the exception of *M. wheeleri* + *M. rubra* (see Supplementary file 2, Fig. S2). The SC phylogeny was topologically similar to the ML phylogeny but was less well supported overall. However, all species within *Manica* received full local posterior probability. One notable topological difference was the placement of one sample of *Manica rubida* (P602). This sample did not amplify well during genomic library preparation; therefore, we recovered fewer UCE loci after enrichment and sequencing for this specimen in contrast to the remainder of the dataset (see Supplementary file 2, Fig. S3; Supplementary Table 4).

In both cases, our backbone phylogeny results show strong support for a set of relationships within *Manica* which has not yet been proposed: the northern Nearctic species *M. hunteri* was recovered as the sister species to the western Palearctic *M. rubida;* the eastern Palearctic *M. yessensis* is strongly supported as sister to the Nearctic *M. invidia* + *M. bradleyi.* These results contradict the assumptions made about *M. invidia* by Wheeler & Wheeler (1970), who proposed that *M. invidia* was the most “ancestral” of the genus due to its generalized morphology, large geographic range, and tolerance of a broad spectrum of habitats. While the term “ancestral” is not used in contemporary systematics when discussing extant species (after all, an extant species being ancestral to other extant species defies logic), at the time the Wheelers were working the term “ancestral” or “primitive” when applied to extant species usually referred to an isolated species that was proposed as being sister to the remainder of the focal group, and to species exhibiting morphological or behavioral traits that were thought to be plesiomorphic. In any case, we did not find this relationship in our study. Furthermore, whereas Zharkov et al. (2022) appear to elect *Manica yessensis* for the extant species with the most “ancestral” characters, we do not find support for *M. yessensis* being the sister lineage to all other extant *Manica* species. Instead, we find a that *Manica rubida* + *hunteri* form the sister group to a clade where *M. yessensis* is sister to *M. invidia* + *M. bradleyi*.

Regarding *Myrmica*, our sampling was relatively sparse. However, here too we found relationships that had not been recognized previously: for example, *Myrmica incompleta* was found to form a clade with *M. punctiventris*, whereas it formed a grade in the most recent phylogeny of *Myrmica* (Jansen et al. 2010). Our backbone phylogeny also included several species that were not present in Jansen et al.’s (2010) study: the Palearctic *Myrmica specioides* was recovered as sister to a large clade composed of Nearctic taxa and *M. mexicana*.

### Morphology

To estimate the phylogenetic position of the Baltic amber fossil †*Manica andrannae,* we gathered morphometric data from all extant *Manica* and *Myrmica* specimens used in this study (*n* = 62; Supplementary table 5) and plotted the means of the morphometric data and indices against morphometric data for †*Manica andrannae* derived from Zharkov et al. (2022). Overall, we found that morphometric measurements differed significantly for *Manica* vs. *Myrmica*, with most of the measurements and indices (36/38) being recovered as significantly different (see Supplementary file 3). Of these, †*Manica andrannae* fell outside of the distributions of *Myrmica* in more than a third (14/38). Crucially, †*Manica andrannae* fell outside of the distributions of all extant *Manica* in ESL, PPL, PrdL, CI, SI2, FLI, OI2, ESLI, ESDI, and PRI, suggesting that †*M. andrannae* is not in the *Manica* crown group, *i.e.* not closely related to any extant *Manica* species.

### *Manica* Fossil Placement and Divergence Dating

To place †*Manica andrannae* within the context of overall Myrmicini evolution, we performed a total-evidence analysis of a combined reduced UCE and morphology dataset. Under ML analysis, the reduced UCE dataset produced a tree very similar in topology and support to the full dataset, apart from the topology of *Myrmica wheeleri* + *Myrmica rubra* and the bootstrap support of *Manica rubida* + *Manica hunteri*. The results of the ML analysis concur with the results of our molecules-only MRBAYES analysis (see Supplementary file 2, Fig. S4).

Our morphology-only MRBAYES analysis recovered *Manica* and *Myrmica* as reciprocally monophyletic and with high support, although the sub-generic relationships were poorly supported. More importantly, we inferred †*Manica andrannae* to be sister to all extant *Manica* with reasonably high support for a morphology-only dataset (0.74 posterior probability; see Supplementary file 2, Fig. S5). Our combined morphology and UCE non-clock MRBAYES analysis recovered the same relationship regarding †*Manica andrannae* as above, with the statistical support of the crown *Manica* weakened (0.86 posterior probability; see Supplementary file 2, Fig. S6). However, when the dataset was inferred with a molecular clock, the same topology is recovered with improved overall support for the topology (see Fig. 2; Supplementary file 2, Fig. S7; see Table 1 for divergence data). When we constrained †*Manica andrannae* to be sister to *M. yessensis,* node ages increased an average of 19.8 Ma over the entire tree, and uncertainty around node ages increased by nearly three-fold on average (see Fig. 2 inset; Supplementary file 2, Fig. S8).

**Figure 2.**
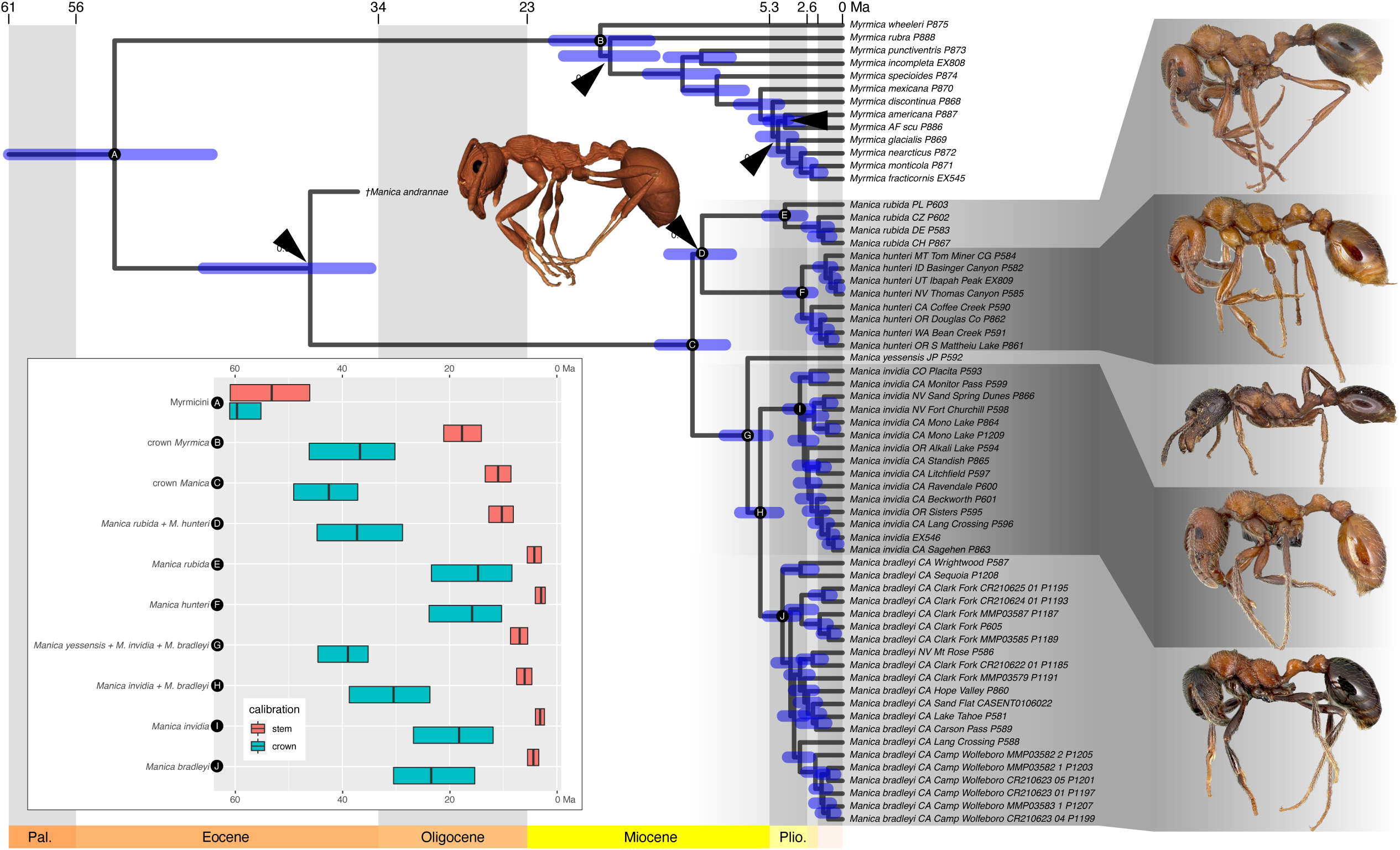
Results of the total evidence analysis, placing the fossil *†Manica andrannae* within the ant tribe Myrmicini. Node bars indicate 95% highest posterior density; all nodes received full posterior probability support (1.0), except where indicated. Inset: comparison of divergence times at specified nodes (A–J) with *†M. andrannae* constrained to the stem of *Manica* (red bars) vs. constrained to be sister to *M. yessensis* (blue bars); bars indicate 95% highest posterior density.

**Table 1.**
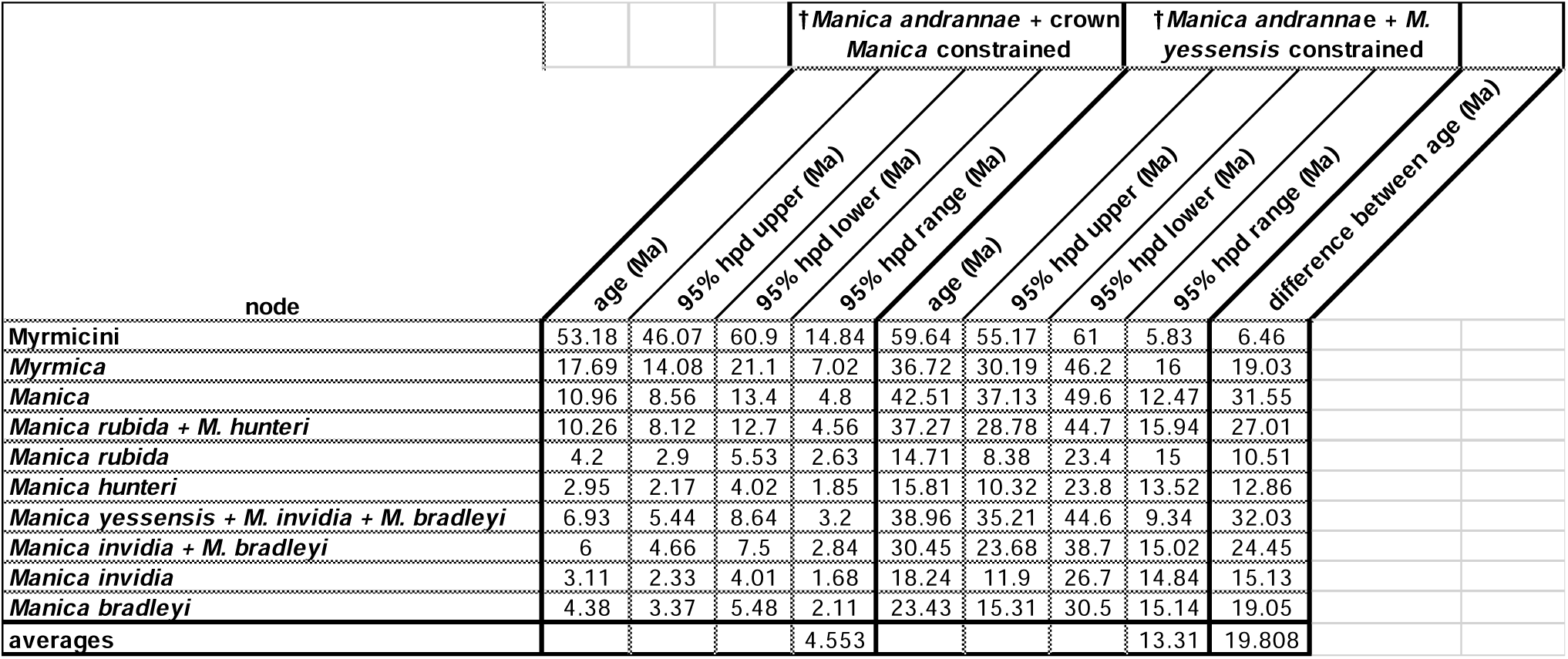
Results of divergence dating analyses, comparing the fossil ant *†Manica andrannae* constrained to the stem of *Manica* vs. *†M. andrannae* constrained to be sister to *Manica yessensis*.

### *Manica* Fossil Placement Effects on Myrmicinae Divergence Dating

For a broader perspective on the effect of fossil placement on divergence dates in the subfamily Myrmicinae, we performed fossil placement experiments using MCMCTREE. This experiment was inspired by the classic study by Ronquist et al. (2012), in which the effects of point-calibration vs. total evidence analysis of fossil placement were compared extensively.

After alignment, trimming, and filtering of the Myrmicinae dataset, the resulting 176 taxon matrix was 1.4 Mb long and contained 2,255 UCE loci, with 15.8% gaps or missing data (see Supplementary Table 4). The ML phylogeny was statistically well supported, with most relationships receiving full bootstrap support (see Supplementary file 2, Fig. S9). The SC phylogeny was less well supported overall and diverged topologically from the ML phylogeny in two notable areas: [Myrmeciinae + [Aneuritinae + Dolichoderinae]], and [Ectatomminae + Formicinae] were recovered, contrary to the findings of any previous study (see Supplementary file 2, Fig. S10). However, the relationships of the myrmicine tribes were identical to Branstetter et al. (2017a).

We performed two divergence dating analyses with MCMCTREE, in which we compare alternative hypotheses on the placement of †*Manica andrannae:* crown (within the clade of extant species) vs. stem (arising from the branch subtending the crown group). We then compared the age distributions around each node in the divergence (Fig. 3, left; Supplementary file 2, Fig. S11 & S12), and found that placing †*Manica andrannae* in the *Manica* crown group not only had a large effect on the age of *Manica* and the crown Myrmicini, but also had widespread effects on the age distributions throughout the formicoid clade, increasing the age of each node by 3 Ma on average (range: 0.1–20.2 Ma). While these two analyses used point-calibrations (as opposed to a total-evidence analysis), the decision to place †*Manica andrannae* as a stem fossil and to demonstrate the consequences of arbitrary placement was informed by the total-evidence analysis of the Myrmicini above. Our results indicate that improper fossil placement can have large effects not only on the divergence estimates of the parent taxon, but also on the broader phylogeny. The stem vs. crown fossil issue is widely acknowledged, and in some cases directly addressed (Boudinot et al. 2022a & b). We strongly suggest that model-based fossil placement approaches should be employed where possible to improve divergence-dating estimates. These approaches have existed for over a decade now (Pyron 2011; Ronquist et al. 2012) but the widespread use of these methods has yet to be fully adopted.

**Figure 3.**
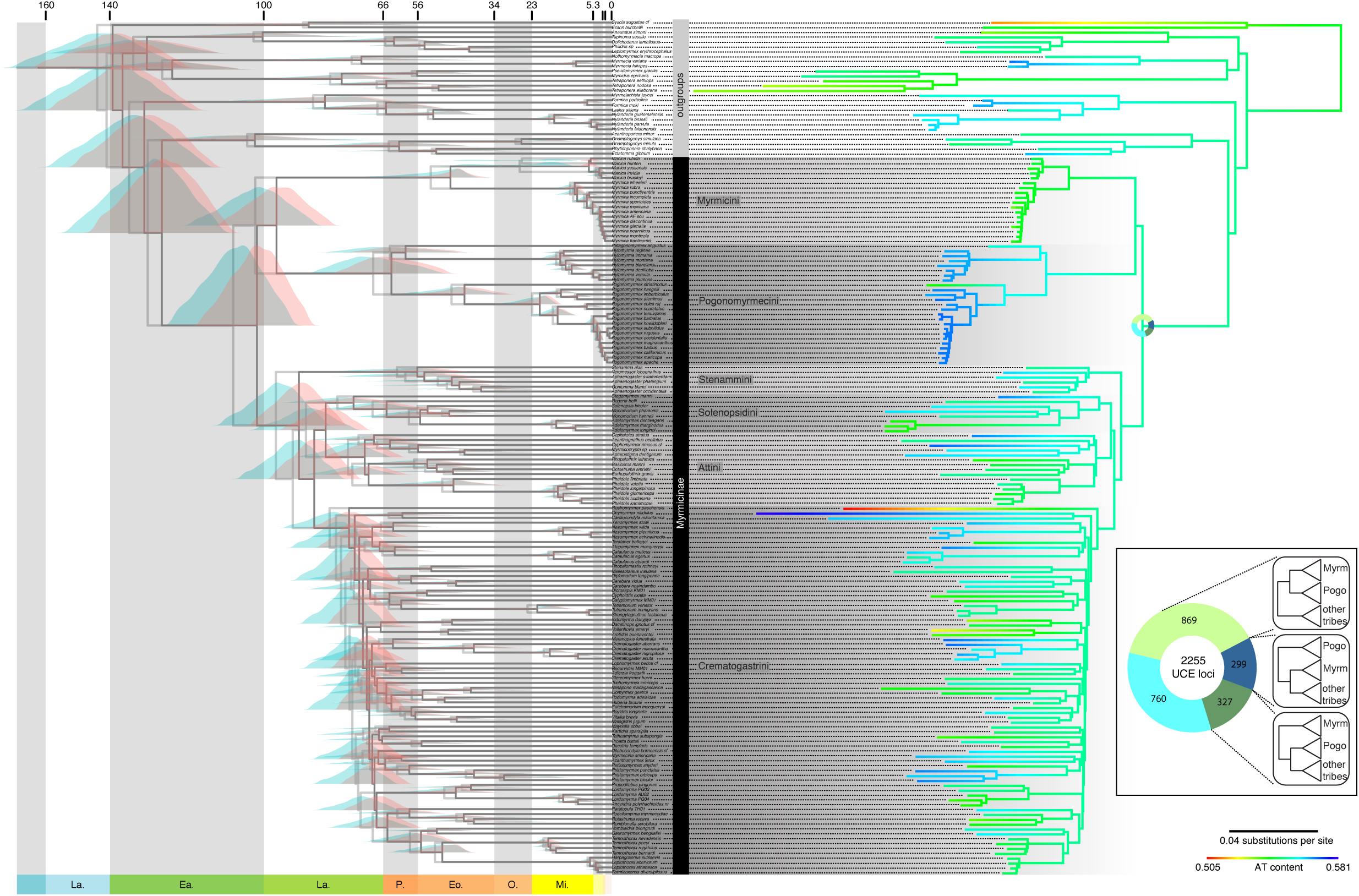
Divergence dating and data characteristics of the ant subfamily Myrmicinae. **A** Effects of *Manica* fossil placement on divergence dating in the Myrmicinae. Purple distributions and dark grey branches correspond to *†Manica andrannae* constrained to the stem of *Manica;* red distributions and light grey branches correspond to *†M. andrannae* constrained to be sister to *M. yessensis.* **B** Phylogram of the Myrmicinae with branches colored to indicate AT content. Inset: Results of the concordance factor analysis, with each slice of the chart indicating number of UCE loci recovering the corresponding topology.

### Tribal Topology

#### Topology tests

The results of the topology tests strongly support the Branstetter et al. (2017a) topology, Myrmicinae + Pogonomyrmecini (Fig. 1b; see Supplementary file 2, Fig. S13), across all tests (see Table 2). The Romiguier et al. (2022) topology (Fig. 1c; see Supplementary file 2, Fig. S14) was only slightly more supported than the Ward et al. (2015) topology (Fig. 1a; see Supplementary file 2, Fig. S15).

**Table 2.** Results of topology tests comparing three distinct tribal topologies for the ant subfamily Myrmicinae.

| Tree | logL | deltaL | bp-RELL | p-KH | p-SH | p-WKH | p-WSH | c-ELW | p-AU |
| --- | --- | --- | --- | --- | --- | --- | --- | --- | --- |
| Branstetter | -35850211.2 | 0 | 0.498 + | 0.505 + | 1 + | 0.505 + | 0.895 + | 0.5 + | 0.503 + |
| Ward | -35852934.2 | 2723 | 0 - | 0 - | 0 - | 0 - | 0 - | 0 - | 0.000456 - |
| Romiquier | -35852866.1 | 2654.9 | 0 - | 0 - | 0 - | 0 - | 0 - | 0 - | 0.00123 - |

#### Gene tree/species tree discordance

Gene concordance analysis revealed that a large proportion of gene trees (869/2255 UCE loci) were in concordance with the Branstetter et al. (2017a) topology (see Fig. 3, inset). The Romiguier et al. (2022) and Ward et al. (2015) topologies were much less represented in analysis of gene trees, with proportions of 299/2255 and 327/2255 UCE loci, respectively.

#### Model adequacy

We found broad variation in model adequacy across test statistics (see Supplementary table 6). Among loci that were concordant with our topologies of interest, we found the most disparity in modeling of GC variance (Ward et al. 2015 topology; Supplementary table 7), mean GC (Branstetter et al. 2017a topology; Supplementary table 8), and maximum pairwise distance (Romiguier et al. 2022 topology; Supplementary table 9). We interpret these test statistics following Höhna et al. (2018): GC variance tests how compositionally heterogeneous an alignment is: significant p values indicate alignments that violate the assumption of homogeneity; mean GC content is a measure of how adequately nucleotide composition of an alignment is modeled; maximum pairwise distance is sensitive to how adequately site rate variation is modeled. For our p4 analyses, we found that 6376/6765 data partitions passed the chi-squared test of homogeneity at a threshold of *p* = 0.05, but only 546/6765 data partitions were compositionally homogenous under the simulation-based test.

Support for the Branstetter et al. (2017a) topology was complete and uniform across unpartitioned and partitioned supermatrix analyses, regardless of which statistic was used to construct matrices (Table 3). However, the topological results of the supertree analyses varied broadly, recovering the Romiguier et al. (2022) topology as an alternate to the Branstetter et al. (2017a) topology in the full, chi-squared, and GC variance datasets, but only when the uncollapsed supertree was used as input. The Ward et al. topology was not observed in any results. In our analyses, we see what appears to be the cumulative effects of two factors which affect the tribal topology of the Myrmicinae: compositional heterogeneity and systematic poor statistical support in the supertree analyses.

**Table 3.** Results of the sensitivity analyses, comparing node support for tribal topologies and *Rostromyrmex* placement across analyses. Myrm = Myrmicini; Pogo = Pogonomyrmecini; Sten = Stenammini; Sole = Solenopsidini; Atti = Attini; Crem = Crematogastrini; Rost = *Rostromyrmex.* Support for supermatrix analyses is in maximum likelihood bootstraps; support for supertree analyses is in posterior probability.

| matrix | method | partitioning | Myrm + Pogo | Myrm + (Sten + Sole + Atti + Crem) | Rost + Crem | Rost + (Sten + Sole + Atti + Crem) | Rost + Myrmicinae | Crem + (Rost + Crem) |
| --- | --- | --- | --- | --- | --- | --- | --- | --- |
| full | supermatrix | unpartitioned | 100 | - | 100 | - | - | - |
| full | supermatrix | SWSC-EN | 100 | - | 100 | - | - | - |
| full | supertree | SWSC-EN | - | 0.96 | - | 1 | - | - |
| full | supertree bs10 | SWSC-EN | 1 | - | 1 | - | - | - |
| max_PD_pass | supermatrix | unpartitioned | 100 | - | - | - | - | 95 |
| max_PD_pass | supermatrix | SWSC-EN | 100 | - | - | - | - | 68 |
| max_PD_pass | supertree | SWSC-EN | 1 | - | 1 | - | - | - |
| max_PD_pass | supertree bs10 | SWSC-EN | 1 | - | 0.99 | - | - | - |
| mean_GC_pass | supermatrix | unpartitioned | 100 | - | 100 | - | - | - |
| mean_GC_pass | supermatrix | SWSC-EN | 100 | - | 100 | - | - | - |
| mean_GC_pass | supertree | SWSC-EN | 0.17 | - | - | - | 1 | - |
| mean_GC_pass | supertree bs10 | SWSC-EN | 1 | - | 1 | - | - | - |
| p4_corrected | supermatrix | unpartitioned | 100 | - | 100 | - | - | - |
| p4_corrected | supermatrix | SWSC-EN | 100 | - | 100 | - | - | - |
| p4_corrected | supertree | SWSC-EN | 0.57 | - | - | 0.66 | - | - |
| p4_corrected | supertree bs10 | SWSC-EN | 0.98 | - | - | - | 0.99 | - |
| p4_uncorrected | supermatrix | unpartitioned | 100 | - | 100 | - | - | - |
| p4_uncorrected | supermatrix | SWSC-EN | 100 | - | 100 | - | - | - |
| p4_uncorrected | supertree | SWSC-EN | - | 0.97 | - | 1 | - | - |
| p4_uncorrected | supertree bs10 | SWSC-EN | 1 | - | 1 | - | - | - |
| var_GC_pass | supermatrix | unpartitioned | 100 | - | 100 | - | - | - |
| var_GC_pass | supermatrix | SWSC-EN | 100 | - | 100 | - | - | - |
| var_GC_pass | supertree | SWSC-EN | - | 0.8 | - | 1 | - | - |
| var_GC_pass | supertree bs10 | SWSC-EN | 1 | - | 1 | - | - | - |

It is worth noting that other than the tribal topology of the Myrmicinae, *Rostromyrmex pasohenis* was observed to have an unstable position among analyses, recovered variously as sister to Crematogastrini, nested within Crematogastrini, sister to all Myrmicinae excluding Myrmicini + Pogonomyrmecini, or sister to all Myrmicinae. Because it is outside of the scope of this study, we did not explore factors that affect the placement of *Rostromyrmex* directly, but due to its high GC content in contrast to the overall dataset (Fig. 3, right), compositional heterogeneity is a likely candidate.

Considering these results, we find that one aspect of the previous myrmicine tribal scheme is supported, in which Myrmicini and Pogonomyrmecini were united under Myrmicini Lepeletier de Saint-Fargeau 1835 (except for *Eutetramorium*, *Huberia*, and *Secostruma*, all of which are in the Crematogastrini). Ward et al. (2015) separated Pogonomyrmecini from Myrmicini based partly on the results of phylogenetic topology, but also because these tribes are easily diagnosable. Here, we retain the six-tribe scheme proposed by Ward et al. (2015) due primarily to the distinct morphology and geographical distributions of the Myrmicini and Pogonomyrmecini.

### Biogeography

We found strong support for our model with potential for overwater dispersal using Bayes factors, calculated from stepping stone analysis (BF_10_=40; see Supplementary file 2, Fig. S18 for results of the divergence dating analysis; Fig. S19 & S20 for results of the ‘dispersal’ and ‘no dispersal’ models). Our estimate of the biogeographical history of the Pogonomyrmecini + Myrmicini recovered the common ancestor of both tribes emerging on the South American continent during the late-Cretaceous (∼100 Ma; see Fig. 4). While our results are ambiguous regarding the geographic origin for the Myrmicini, the crown *Myrmica* were reconstructed as having a western Nearctic origin with modest support.

**Figure 4.**
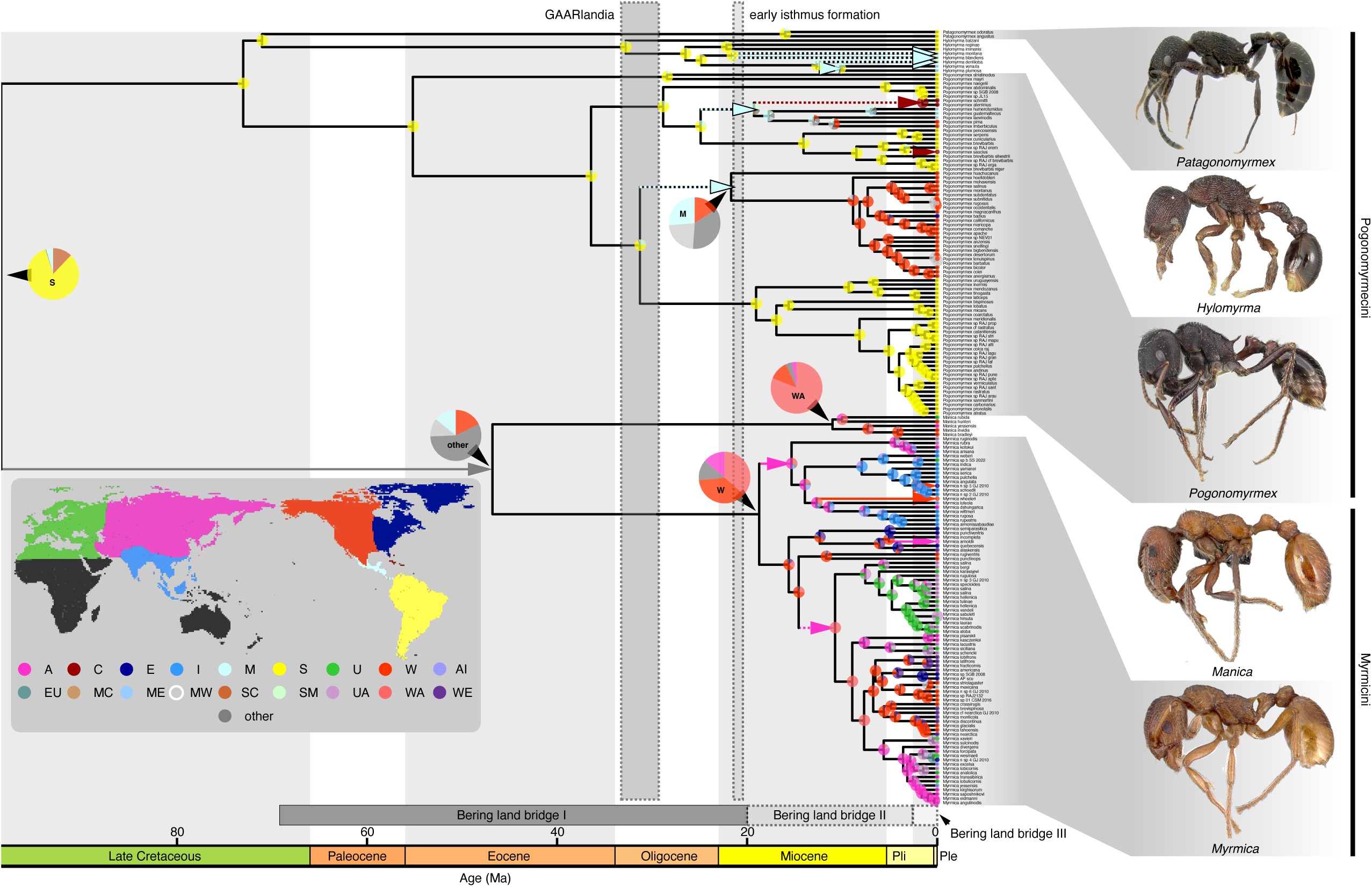
Historical biogeography of the ant tribes Pogonomyrmecini & Myrmicini. Pie charts indicate distribution posterior probabilities at each node. Inset: map of areas used in this study.

Subsequent dispersal across Beringia to the eastern Palearctic commenced in the early Miocene in *Myrmica*, with some lineages inhabiting a Holarctic range during the late Miocene. The MRCA of *Manica* was inferred to have a Holarctic range as well (with high probability): the current species distributions could be explained by fragmentation of this ancient range as an alternative to dispersal across the Bering land bridge. Strikingly, it is still unclear how the common ancestors of *Myrmica* and *Manica* dispersed to North America and Eurasia from South America due to the long branch subtending the crown Myrmicini (as well as *Manica* and *Myrmica*), suggesting extensive extinction of stem lineages. However, the multiple fossils of *Manica* and *Myrmica* from the Eocene Baltic amber and impression fossils of North America hint at the diversity of the stem lineages, as well as the extensive range that the Myrmicini had achieved in the Northern Hemisphere during the Eocene, prior to the diversification of the crown group (Radchenko et al. 2007; Jessen 2020).

Sanmartín et al. (2010) recognized that dispersal across the Beringia has continued nearly uninterrupted since the late Cretaceous, but falls into three consecutive stages, each of which correspond to specific paleoclimates and associated vegetation types found at high latitudes:

1. Bering land bridge I (70–20 Ma): dispersal of mesophytic forests and their associated warm- temperate groups.
2. Bering land bridge II (20–3 Ma): dispersal of temperate-boreal groups associated with coniferous forests as the climate cooled at higher latitudes
3. Bering land bridge III (3–0 Ma): active intermittently, permitting Arctic biota associated with tundra.

Several recent studies have investigated trans-Beringian dispersal of highly diverse Holarctic ant genera. Notably, Schär et al. (2018) made a first attempt to test whether Holarctic ant species exist in multiple genera that span the Bering strait. Perhaps the most relevant finding of Schär et al. (2018) to our study is that trans-Beringian dispersal occurred repeatedly in both directions throughout the Miocene and Plio-Pleistocene. Additional examples from taxon-specific studies reinforce this finding, with similar events in *Leptothorax* (∼7 Ma; Prebus 2017), *Lasius* (numerous times spanning the Miocene and Plio-Pleistocene; Boudinot et al. 2022), *Formica* (at least 8 times spanning the Miocene and Plio-Pleistocene; Borowiec et al. 2021). These findings concur with our results and highlight the frequent dispersal of temperate-boreal lineages across Beringia throughout the Miocene (Tiffney 1985), consistent with the observation that *Leptothorax*, *Lasius*, *Formica*, and *Myrmica* are all common and speciose in present day northern temperate-boreal forests. In this study, we found only a couple of Plio- Pleistocene dispersal events, but it should be noted that our taxon sampling is not complete. More extensive sampling may reveal additional recent dispersal events.

While we were completing this study, a review of the genus *Manica* was published which described a new species from Sichuan province in China (He et al. 2024). The new species *Manica shanyii* He et al. was found nesting in rotten wood and has 7 masticatory teeth, which is unusual for *Manica* species which typically have 12–14 teeth and nest in soil or under rocks, although *M. rubida* has been documented nesting under bark (Antweb 2024). Future studies into the systematics and biogeography of *Manica* should include this interesting species, although this finding does not contradict our hypothesis of a Holarctic *Manica* common ancestor.

Within the Pogonomyrmecini, we found that the diversification of this tribe mostly took place in the Neotropics, primarily on the South American continent. We found that the proposed formation of the Greater Antilles and Aves Ridge (GAARlandia) 30–33 Ma (Iturralde-Vinent & MacPhee 1999; Iturralde-Vinent 2006) coincides with dispersal of the common ancestor of *Pogonomyrmex huachucana* and *Pogonomyrmex sensu stricto* from South to Central America; however, it should be noted that there has not been a land bridge proposed between the Greater Antilles and mainland Central America. This scenario does not preclude the possibility that this ancestor dispersed across the Central American Seaway (CAS) by long distance flight, rafting, or by island hopping (Oberski 2022; Mera-Rodriguez et al. 2023). We should also note that the extant species on the Greater Antilles were reconstructed as arriving well after the formation of GAARlandia. Bacon et al. (2015) raises an intriguing third possibility: limited terrestrial biotic interchange, associated with Pacific-Atlantic marine biotic separation, appears to have occurred between South and Central America as early as 20 Ma. This may help explain the dispersal of several other groups in the Pogonomyrmecini from South to Central America, including *Hylomyrma* and the common ancestor of *Pogonomyrmex schmitti + aterrimus + guatemalicus + humerotimidus + laevinodis + imberbiculus + pima,* which began diversifying in what is now Central America around 20 Ma. Increasingly, evidence from biogeographical studies support biotic interchanges between South and Central America that predate the commonly accepted ∼3.5 Ma closure of the Isthmus of Panama (Winston et al. 2017 (but see Stange et al. 2018); Branstetter et al.2017b; Price et al. 2022). Puzzlingly, the only known fossil of *Pogonomyrmex* is an impression from the late Eocene (∼34 Ma) Florissant formation in Colorado (Carpenter 1930), which is predates any known North or Central American lineage in our analysis. We also find that *Pogonomyrmex s.s.* began diversifying in North America ∼9 Ma ago, coinciding with the growth of desert ecosystems in southwestern North America and the diversification of the honeypot ants, genus *Myrmecocystus* (van Elst et al. 2021).

## Conclusions

Our study provides insights into potential sources of bias affecting the reconstruction of phylogenetic topology and divergence times within the ants. We find that our current understanding of tribal topology in the Myrmicinae is strongly supported (Branstetter et al. 2017a), but this topology is highly sensitive to the effects of compositional heterogeneity and gene-tree species-tree conflict. Our fossil placement analyses strongly suggest that †*Manica andrannae* is a stem *Manica* species, and that placement of this fossil in the crown group affects not only divergence dates within the tribe Myrmicini, but also has broad implications for divergence times throughout the formicoid clade. Furthermore, we resolve long-standing questions about the biogeographic history of *Mymica* and *Manica*. Our biogeographic reconstructions indicate a South American origin for the Pogonomyrmecini + Myrmicini, with the MRCA of *Myrmica* inhabiting the western Nearctic in the early Miocene prior to repeated dispersal across Beringia throughout the Miocene and Pliocene. The MRCA of *Manica*, on the other hand, was inferred to have a Holarctic range prior to vicariance during the Pliocene, in contrast to previously proposed dispersal hypotheses. Unexpectedly, we found strong support within the

Pogonomyrmecini for three coordinated dispersal events from South to Central America during the early Miocene, which has been previously proposed as an early biotic interchange event prior to the more commonly accepted 3.5 Ma closure of the Isthmus of Panama. It is our hope that this study provides a framework for future inquiries into sources of systematic bias in phylogenic reconstruction and divergence dating, as well as further studies on the complex history of GABI.

## Supplementary Material

Data files and online-only appendices can be found in the Dryad data repository: XXX. Newly generated raw read sequence data have been deposited in the NCBI Sequence Read Archive (SRA): BioProject PRJNA1072056. Morphometric data have been deposited in MorphoBank (http://morphobank.org/permalink/?P5425).

## Acknowledgements

We wish to thank Phil Ward for arranging the loan of specimens from the Bohart Museum of Entomology collection at the University of California, Davis.

