## Supplementary file 1 for "Phylogenomics resolve the systematics and biogeography of the ant tribe Myrmicini and tribal relationships within the hyperdiverse ant subfamily Myrmicinae"

**Supplementary File 1**. Extended methods from Prebus & Rabeling 2024 "Phylogenomics resolve the systematics and biogeography of the ant tribe Myrmicini and tribal relationships within the hyperdiverse ant subfamily Myrmicinae"

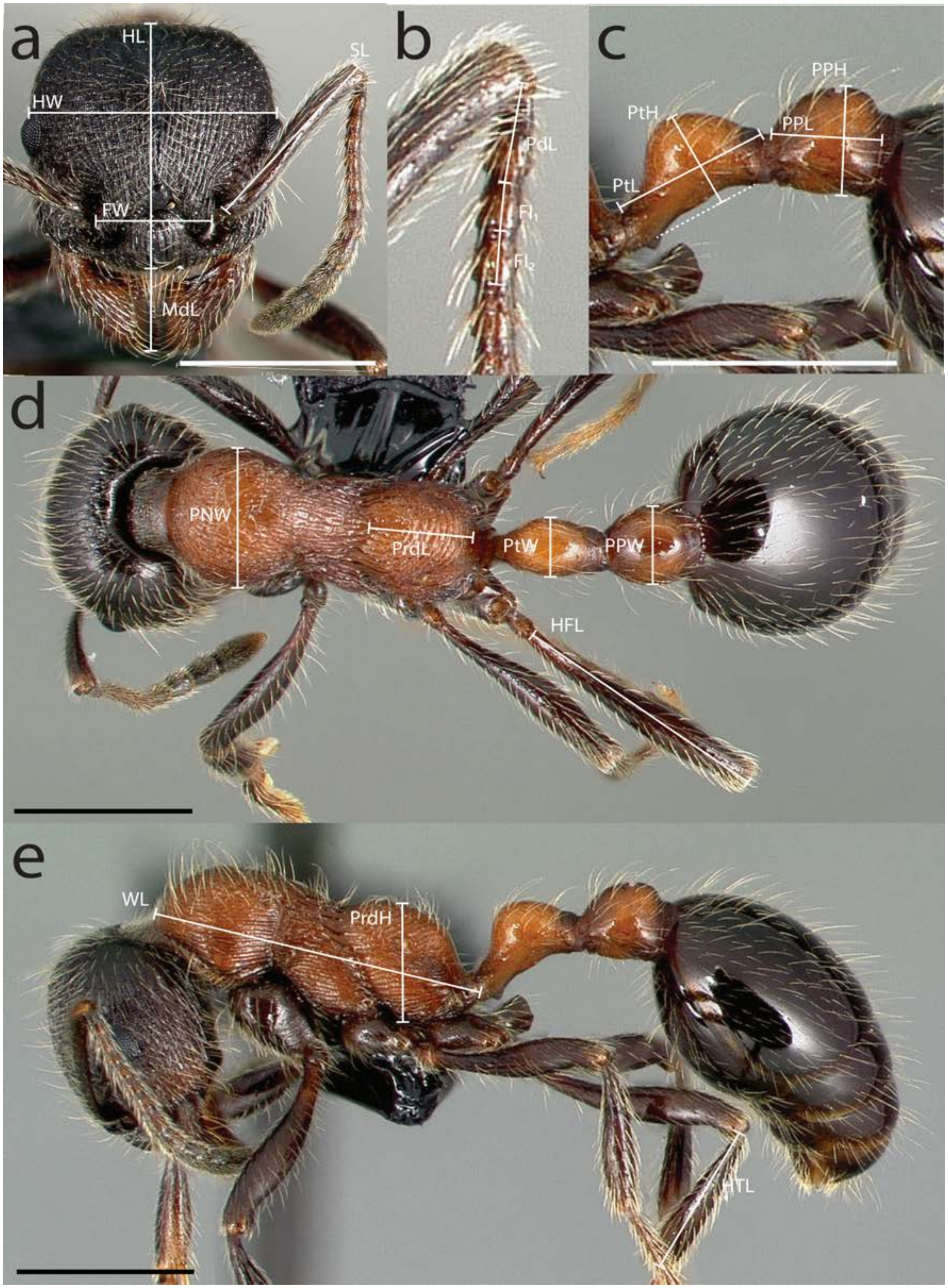


**Fig. S1.** Measurements used in this study demonstrated on *Manica bradleyi* (CASENT0005697) **a** head in full-face view **b** detail of antennae **c** detail of waist segments in profile view **d** dorsal view **e** profile view. Scale bars = 0.2 mm. Photos by April Nobile from www.antweb.org.

**Measurements**

**HL** head length: the head must be carefully tilted to the position, providing the true maximum. If excavations of the posterior margin of the head capsule and/or anterior margin of the clypeus are present, then the measurement is taken from an imaginary line that spans the excavations from the posterior- or anterior-most margins;

**HW** head width: maximum width of the head, measured directly behind the compound eyes;

**FW** frontal lobe width: the maximum width measured between the frontal lobes;

**MdL** mandible length: the length of the mandible, measured from the mandibular apex to the anterior clypeus margin, or to the transverse line connecting the anterior-most points in those taxa where the margin is concave medially;

**SL** scape length: maximum scape length, excluding the basal neck and the articular condyle;

**PdL** antennal pedicellum length;

**$FI_1$** the length of the first flagellomere;

**$FI_2$** the length of the second flagellomere;

**OL** Ocular length: the maximum diameter of the compound eye;

**WL** Weber's length: distance between the caudal most point of propodeal lobe to the inflection point between the pronotal neck and the pronotal declivity;

**PrdH** Propodeum height: the height of the propodeum in profile, measured as the perpendicular distance from the ventral edge to the highest point of the propodeum;

**ESL** Propodeal spine length: maximum length of propodeal tubercles (or spines) in profile, measured along the tubercle/spine from its tip to the deepest point of the propodeal constriction at its base;

**PtL** Petiole length: the length of the petiolar node in profile, measured as the distance from the place of attachment to the propodeum to the place of attachment to the postpetiole;

**PtH** Petiole height: the height of the petiolar node in profile, measured as the perpendicular distance from an imaginary line joining the base of the subpetiolar tooth and the ventral junction of the petiole and postpetiole to the highest point of the petiolar node;

**PPL** Postpetiole length: the length of the postpetiole in profile, measured as the distance from the place of attachment to the petiolar node to the place of attachment to the gaster;

**PPH** Postpetiole height: the height of the postpetiole in profile, measured as the perpendicular distance from the ventral edge to the highest point of the postpetiole;

**PNW** Pronotum width: the maximum width of the pronotum in dorsal view;

**PrdL** Propodeum length: the maximum length of the propodeum in dorsal view;

**ESD** Propodeal spine distance: distance between the tips of propodeal tubercles/spines in dorsal view;

**PtW** Petiole width: the maximum width of the petiolar node in dorsal view;

**PPW** Postpetiole width: the maximum width of the postpetiole in dorsal view;

**HFL** Hind femur length: the maximum length of hind femur, measured in dorsal view;

**HTL** Hind tibia length: the maximum length of hind tibia, measured in anterior view.
