## Supplementary file 2 for "Phylogenomics resolve the systematics and biogeography of the ant tribe Myrmicini and tribal relationships within the hyperdiverse ant subfamily Myrmicinae"

**Supplementary File 2.** Results of phylogenetic analyses and biogeographic history estimations from Prebus & Rabeling 2024 “Phylogenomics resolve the systematics and biogeography of the ant tribe Myrmicini and tribal relationships within the hyperdiverse ant subfamily Myrmicinae”

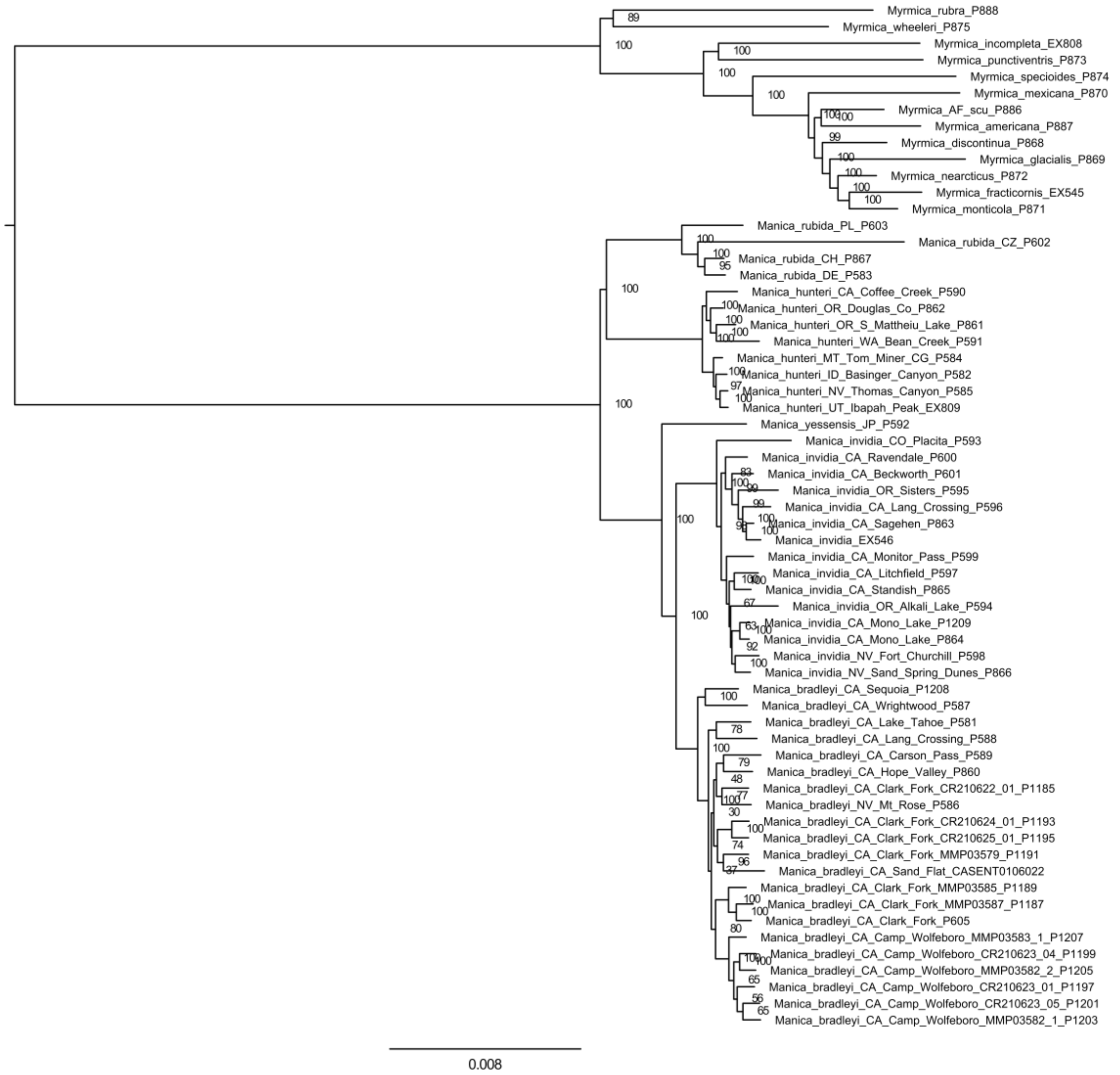


**Figure S2.** Results of maximum likelihood analysis of the ‘Myrmicini UCEs’ dataset with IQTREE2. Statistical support at each node is in ultrafast bootstraps. Branch lengths are in units of substitutions per site.

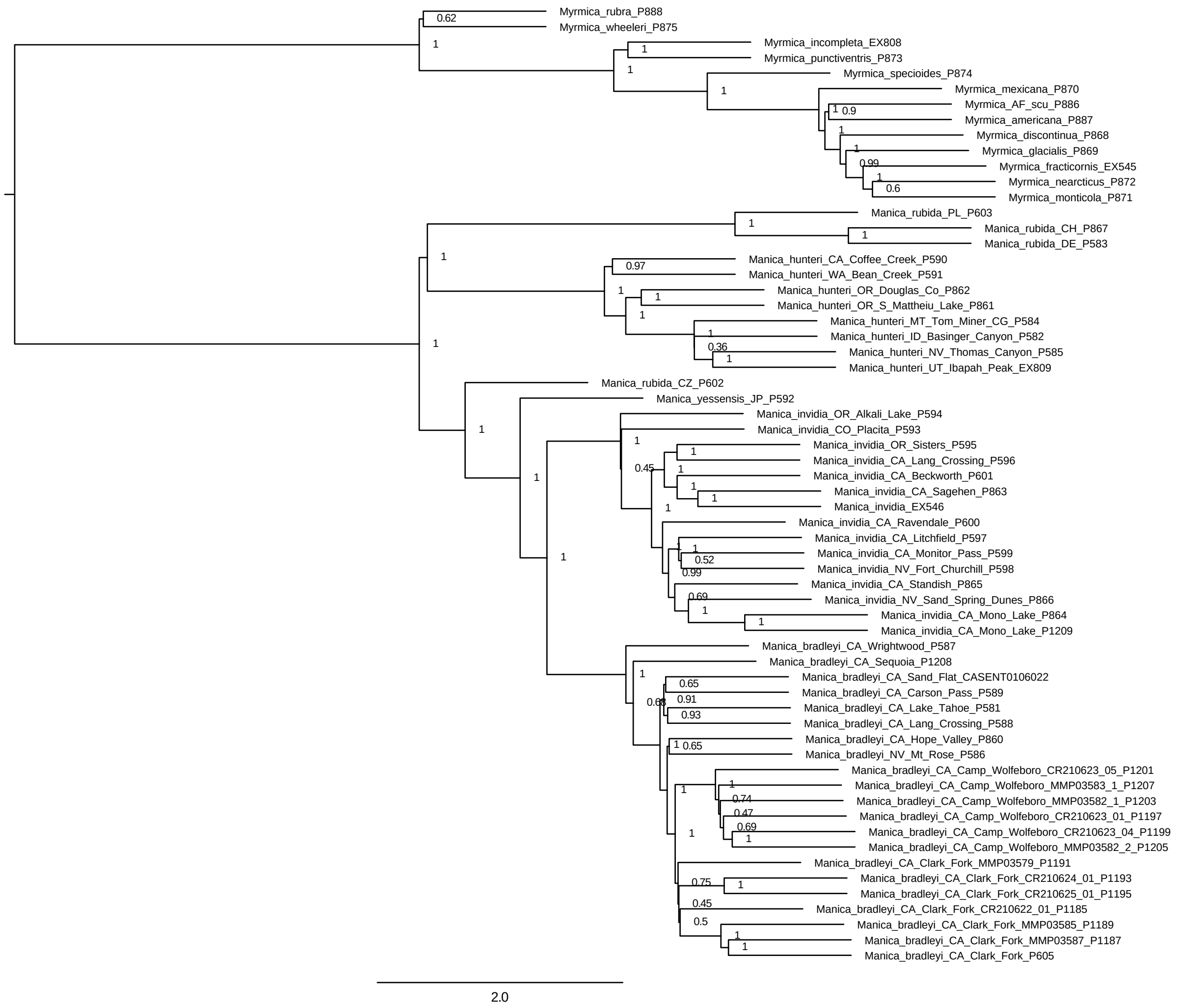


**Figure S3.** Results of summary coalescence analysis of the 'Myrmicini UCEs' dataset with ASTRAL. Statistical support at each node is in local posterior probability. Branch lengths are in coalescent units.

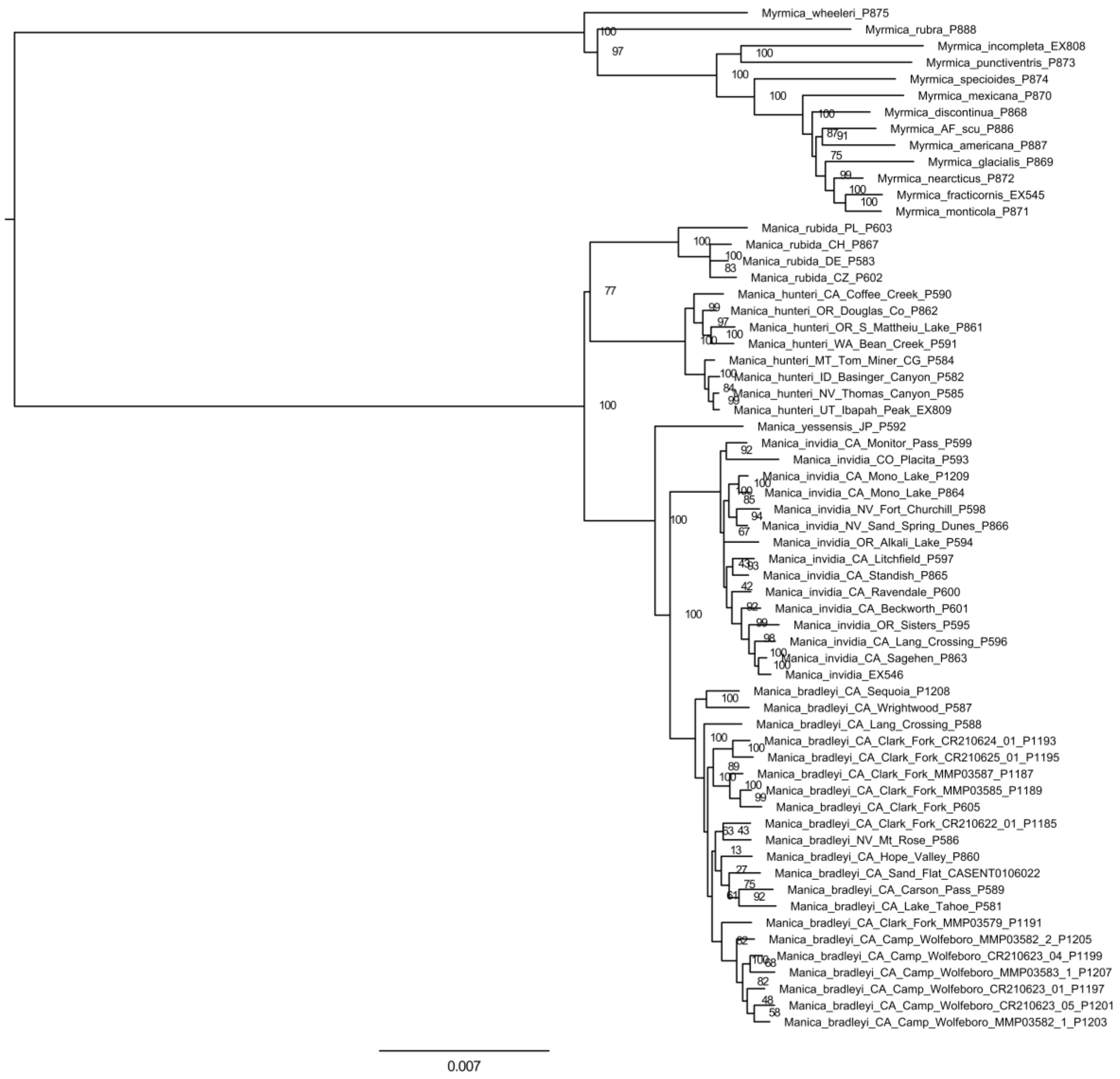


**Figure S4.** Results of maximum likelihood analysis of the ‘reduced Myrmicini UCEs’ dataset with IQTREE2. Statistical support at each node is in ultrafast bootstraps. Branch lengths are in units of substitutions per site.

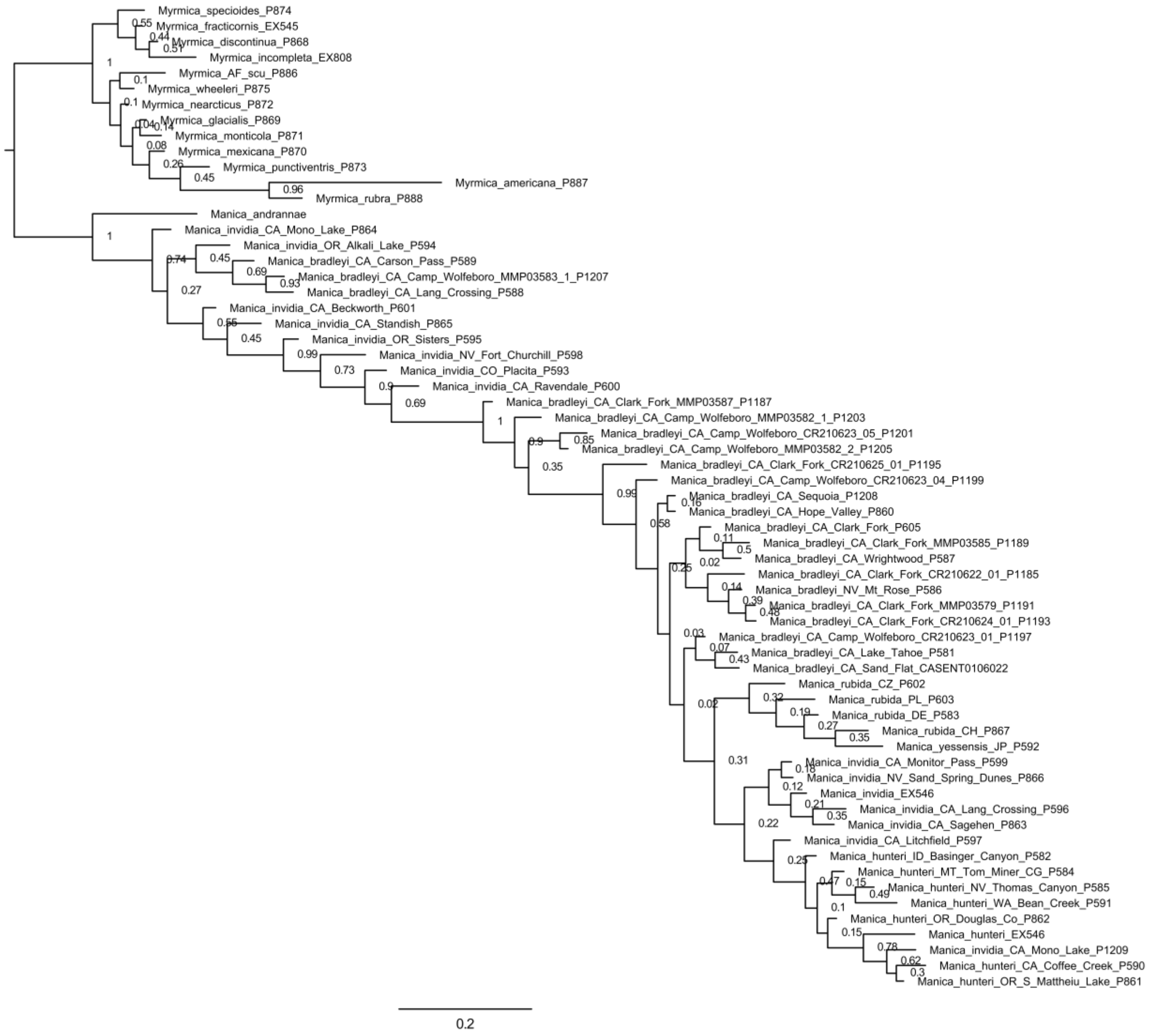


**Figure S5.** Results of Bayesian inference analysis of the ‘Myrmicini morphology’ dataset with MRBAYES. Statistical support at each node is in posterior probability. Branch lengths are in units of substitutions per site.

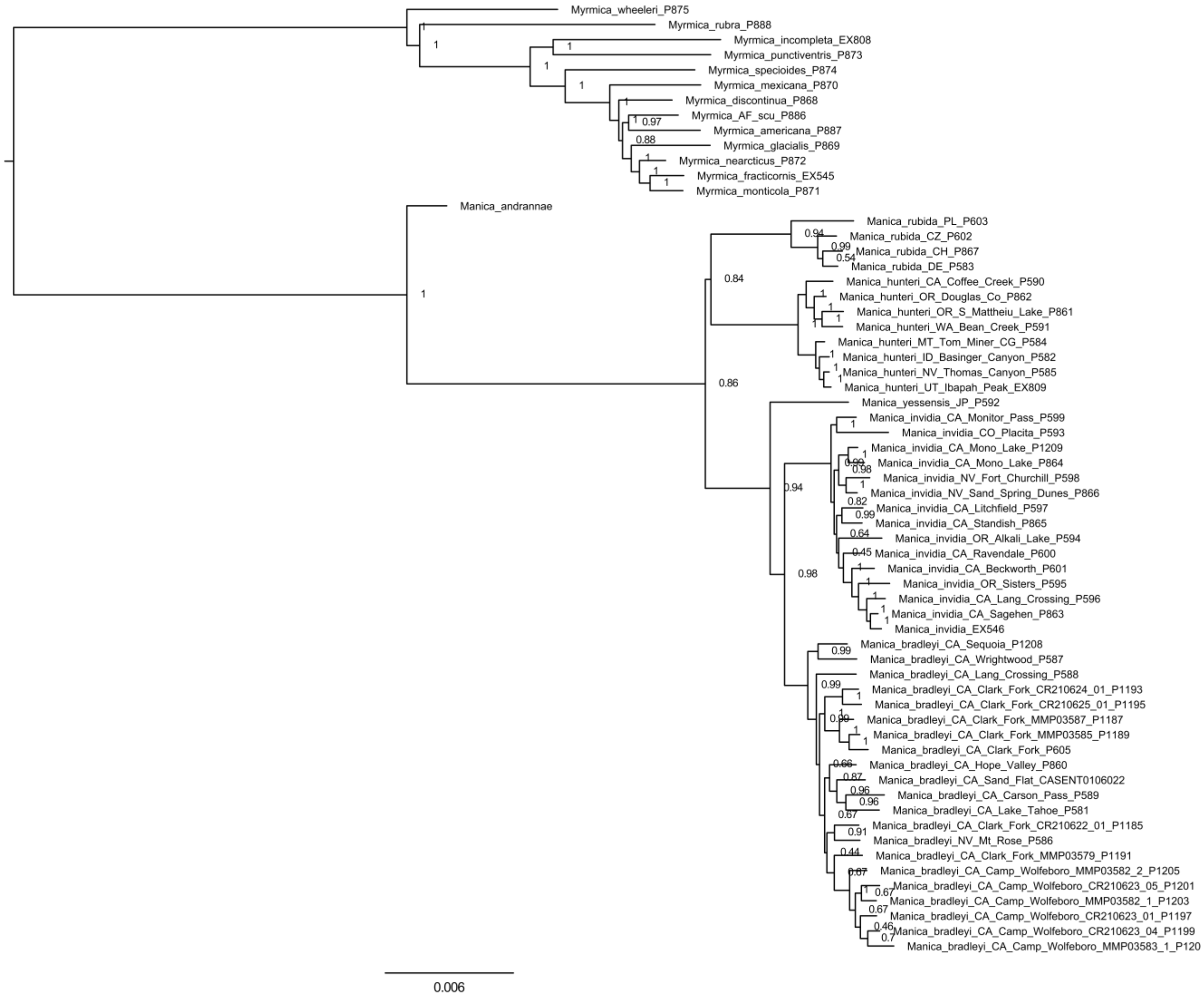


**Figure S6.** Results of Bayesian inference analysis of the 'Myrmicini combined' dataset with MRBAYES. Statistical support at each node is in posterior probability. Branch lengths are in units of substitutions per site.

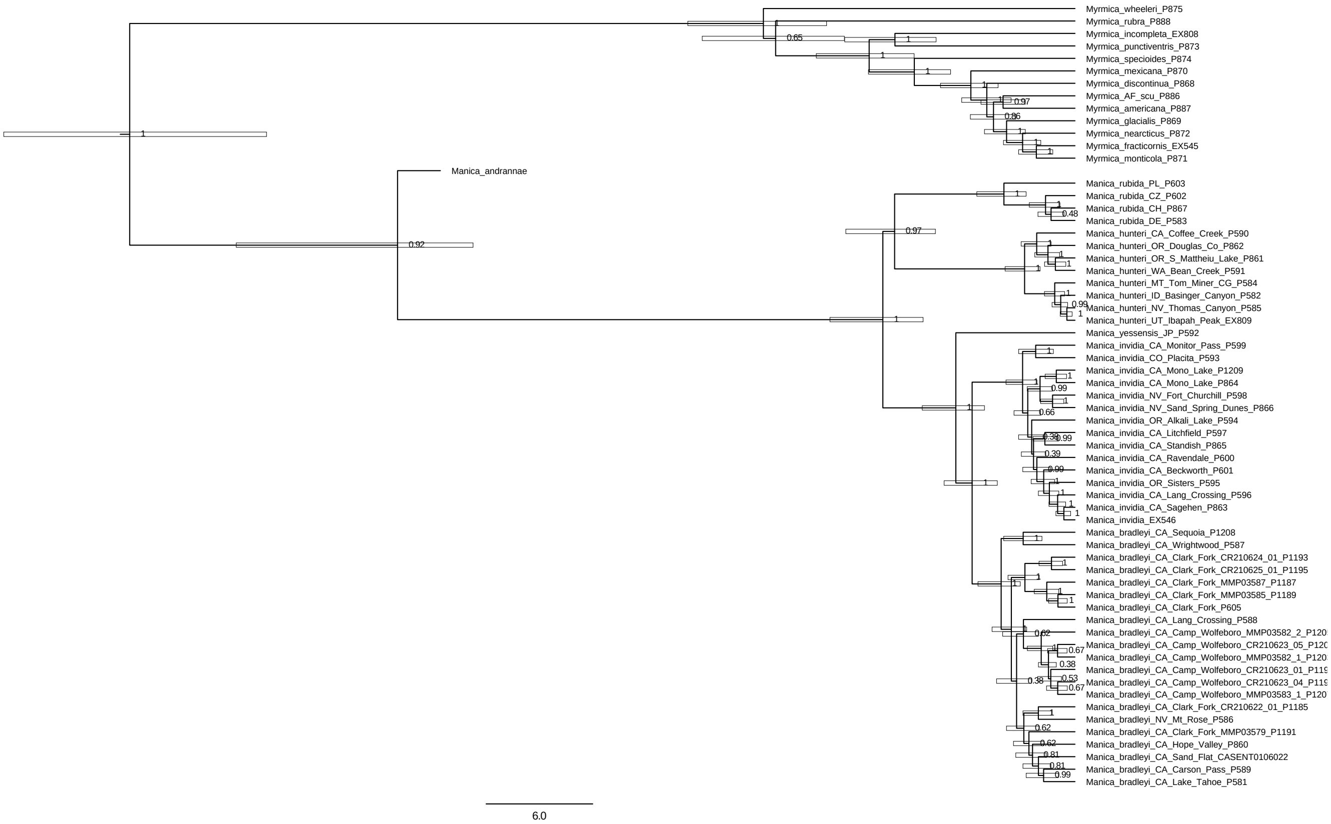


**Figure S7.** Results of the unconstrained Bayesian divergence dating inference analysis of the 'Myrmicini combined' dataset with MRBAYES. Statistical support at each node is in posterior probability. Bars around each node indicate 95% highest posterior density (HPD). Branch lengths are in units of one million years.

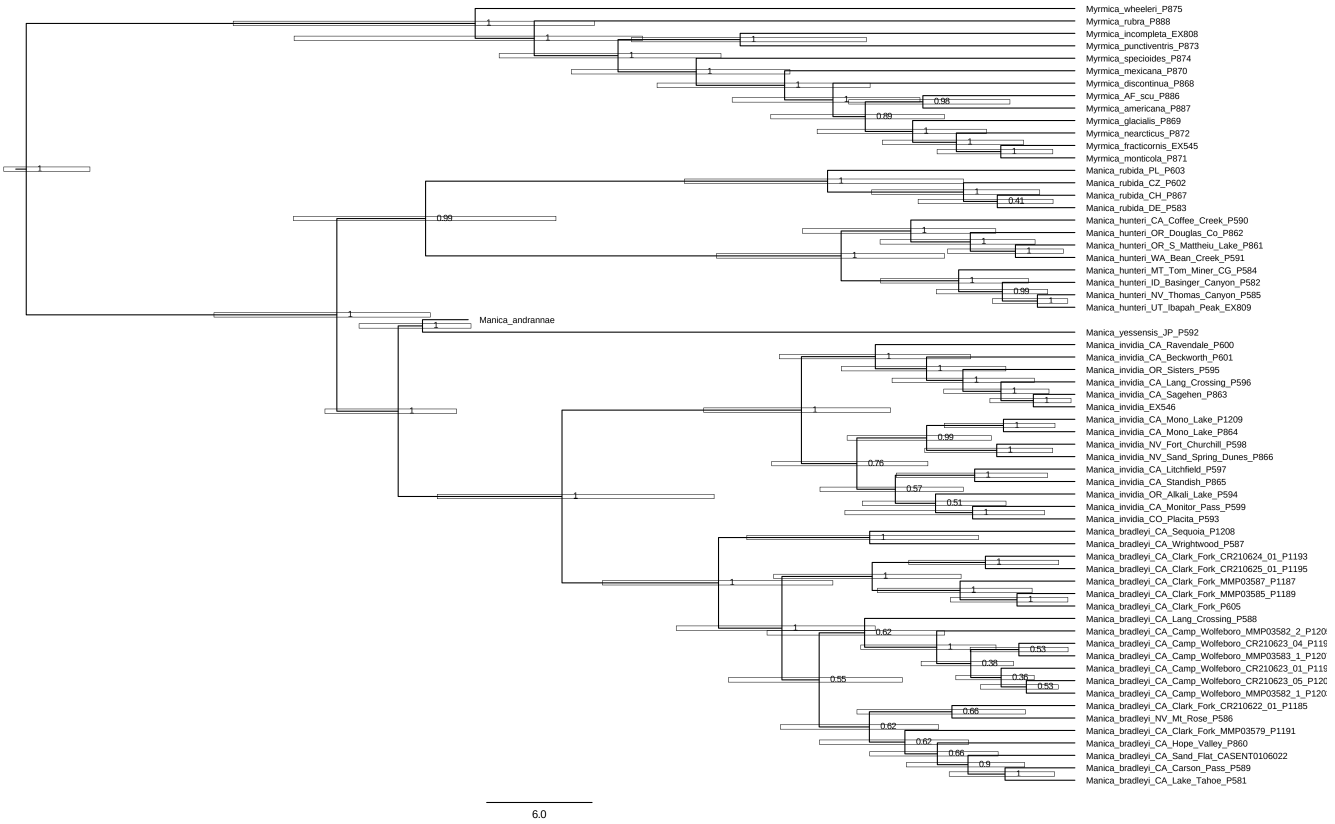


**Figure S8.** Results of the constrained Bayesian divergence dating inference analysis of the ‘Myrmicini combined’ dataset with MRBAYES. Statistical support at each node is in posterior probability. Bars around each node indicate 95% highest posterior density (HPD). Branch lengths are in units of one million years.

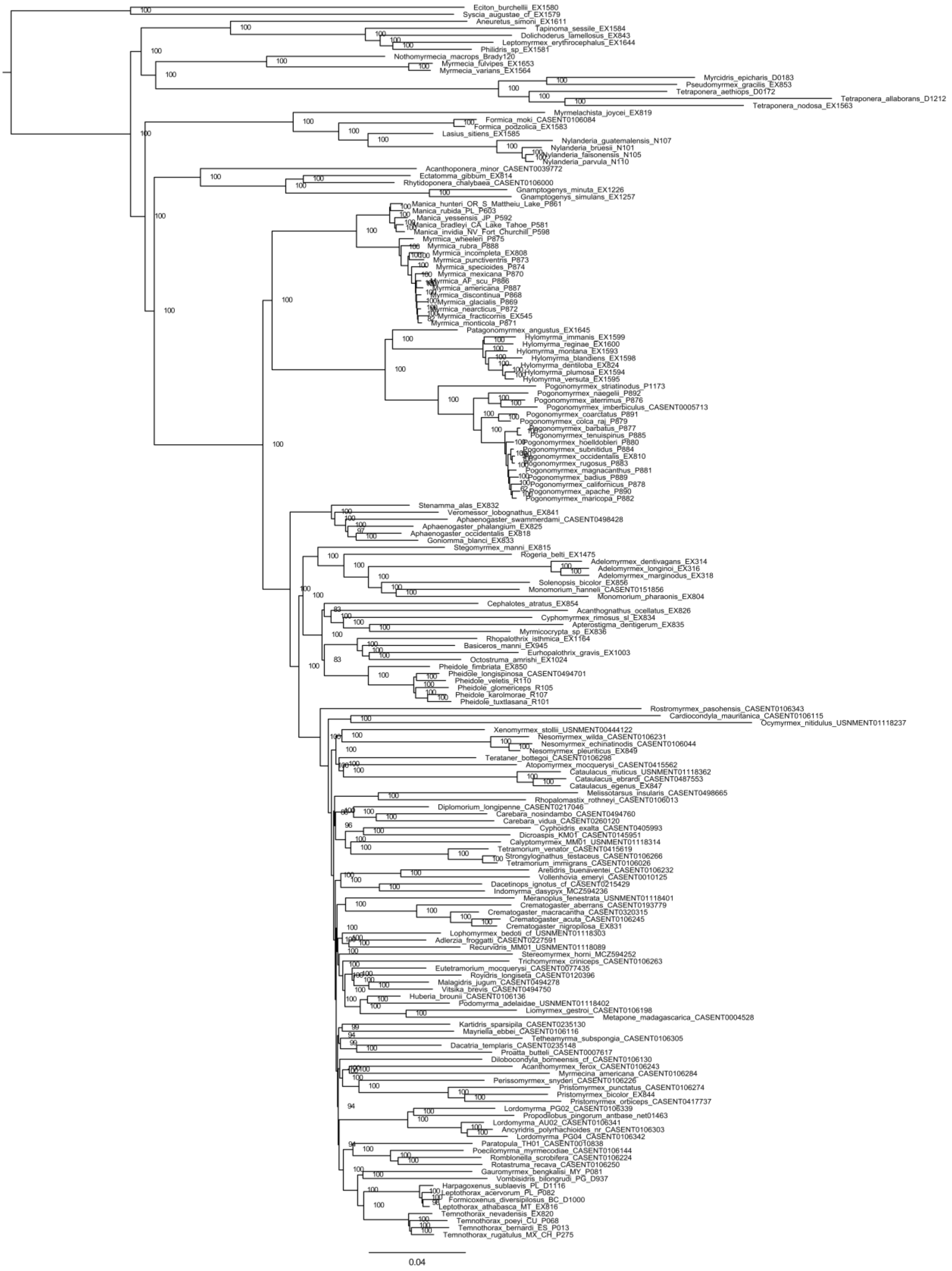


**Figure S9.** Results of maximum likelihood analysis of the ‘Myrmicinae UCEs’ dataset with IQTREE2. Statistical support at each node is in ultrafast bootstraps. Branch lengths are in units of substitutions per site.

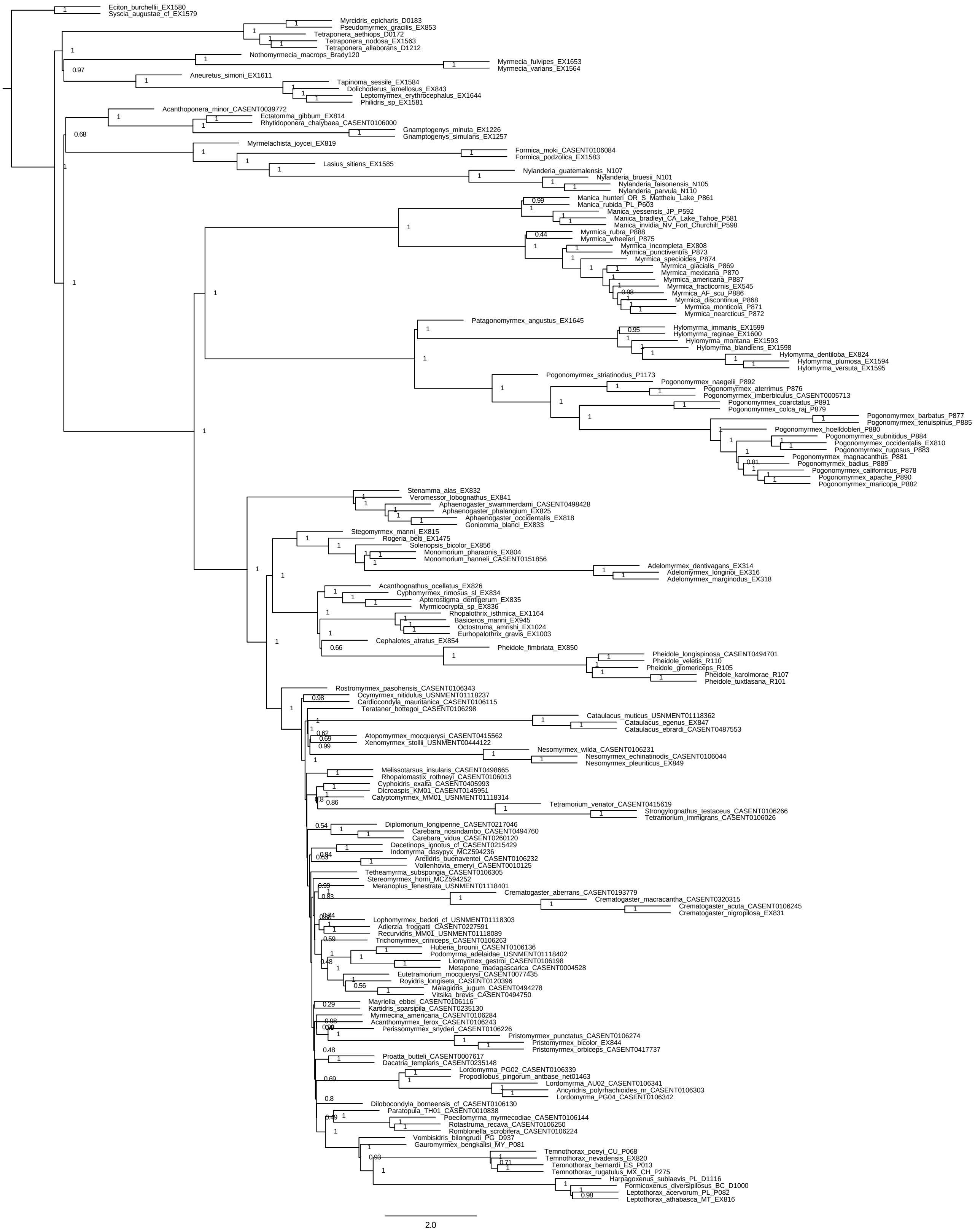


**Figure S10.** Results of summary coalescence analysis of the ‘Myrmicinae UCEs’ dataset with ASTRAL. Statistical support at each node is in local posterior probability. Branch lengths are in coalescent units.

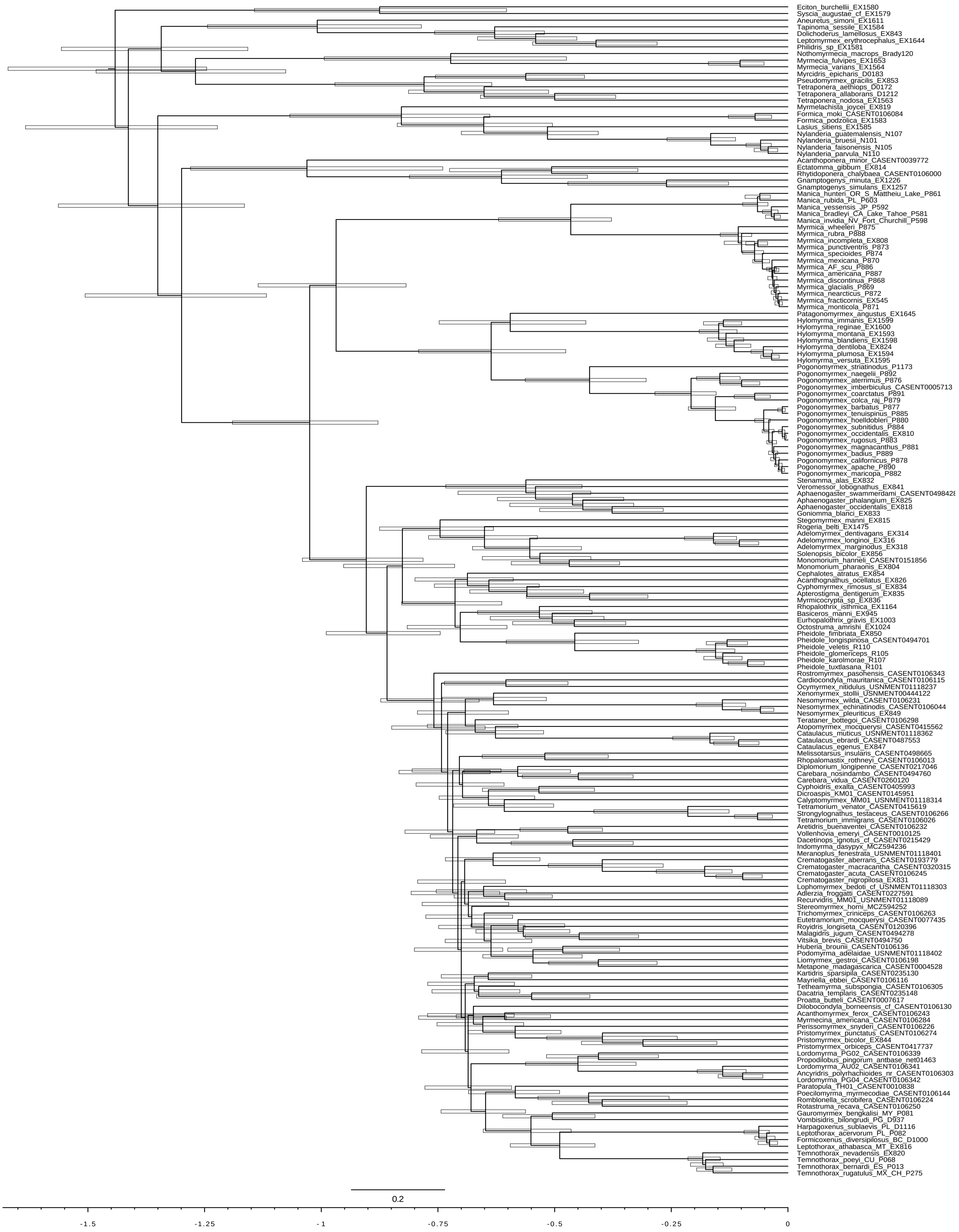


**Figure S11.** Results of divergence dating analysis of the ‘Myrmicinae UCEs’ dataset with MCMCTree assuming a stem position for †*Manica andrannae*. Bars around each node indicate 95% highest posterior density (HPD). Branch lengths are in units of 100 million years.

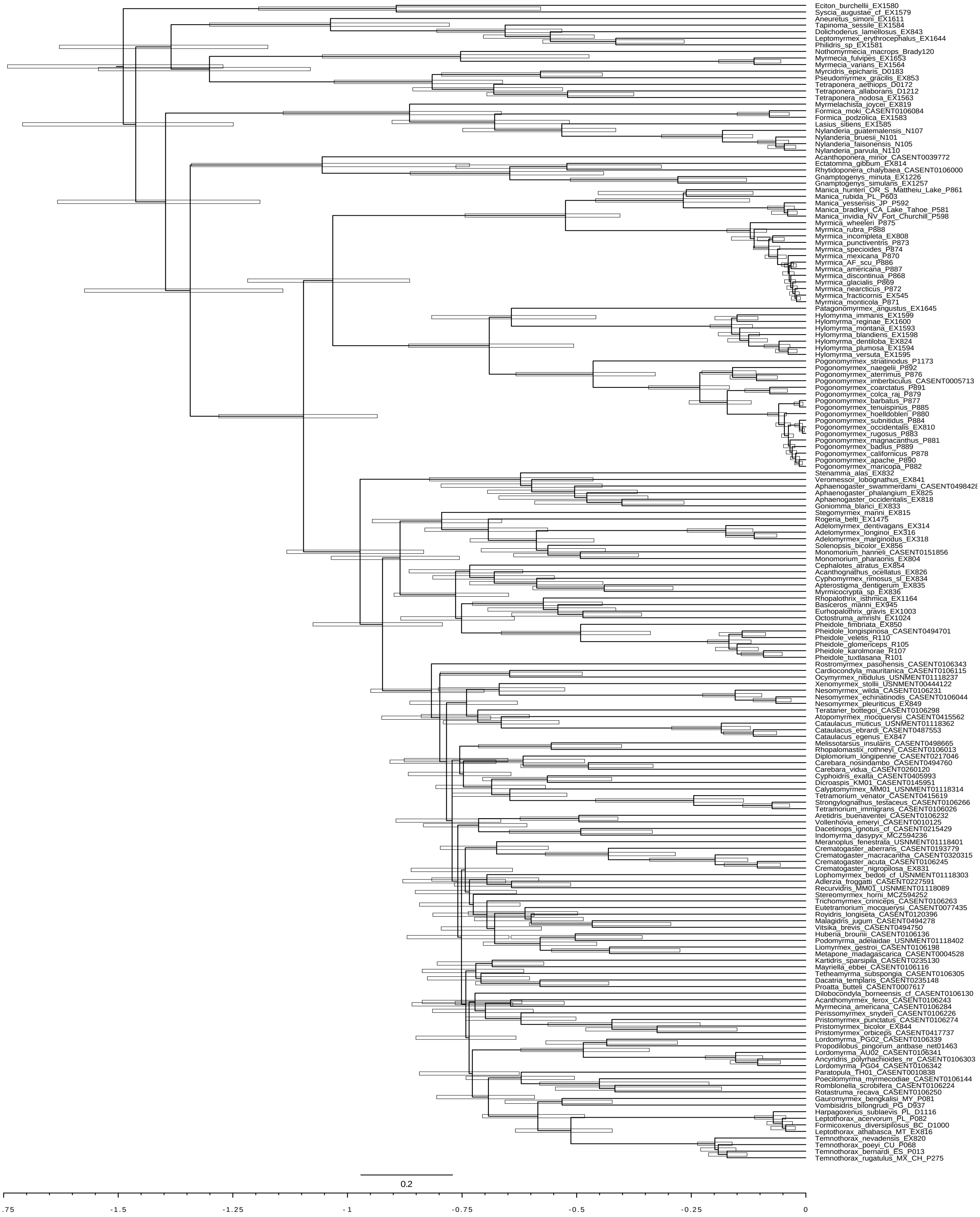


**Figure S12.** Results of divergence dating analysis of the 'Myrmicinae UCEs' dataset with MCMCTree assuming a crown position for †*Manica andrannae*. Bars around each node indicate 95% highest posterior density (HPD). Branch lengths are in units of 100 million years.

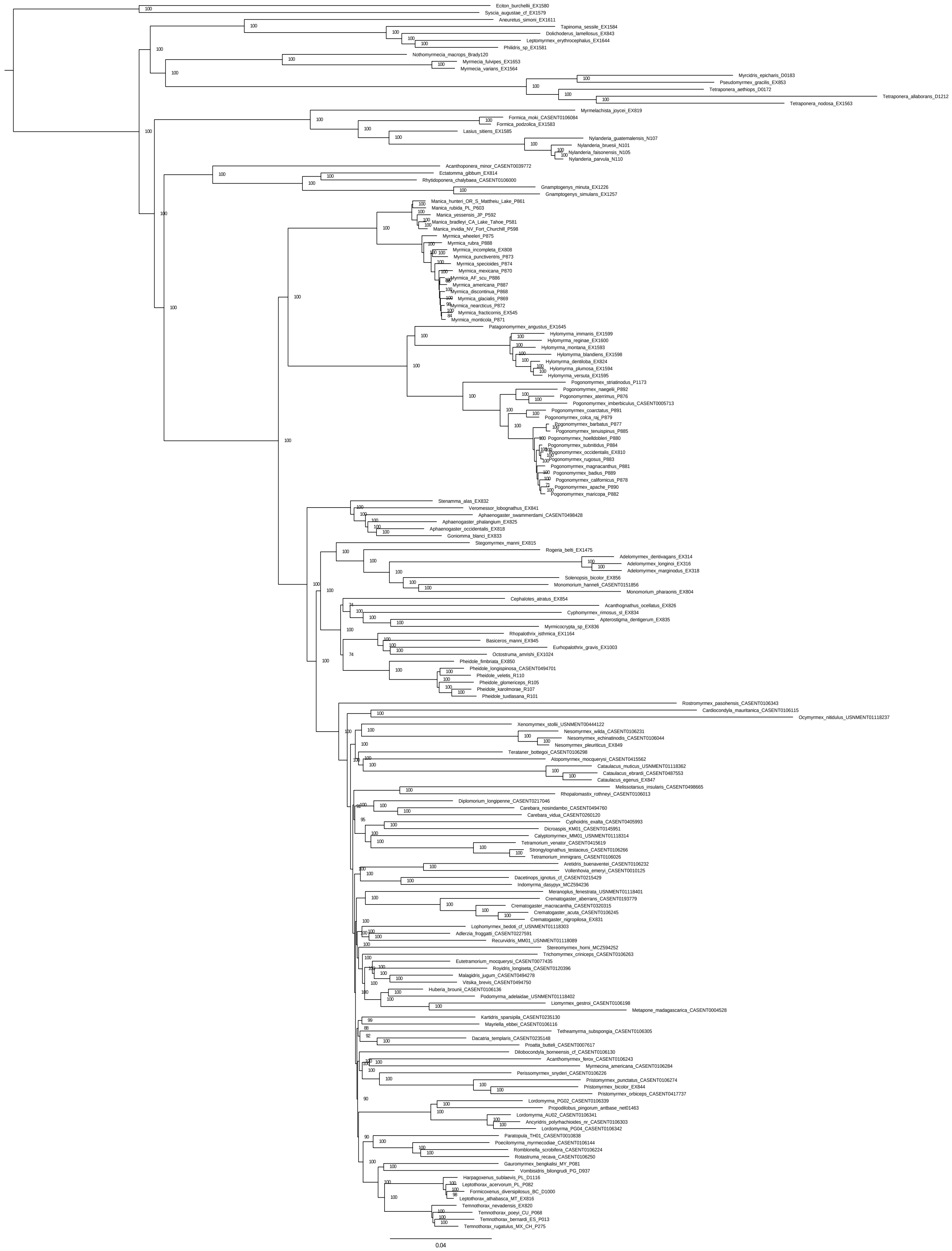


**Figure S13.** Results of maximum likelihood analysis of the ‘Myrmicinae UCEs’ dataset with IQTREE2, constrained to the Branstetter et al. 2017a topology. Statistical support at each node is in ultrafast bootstraps. Branch lengths are in units of substitutions per site.

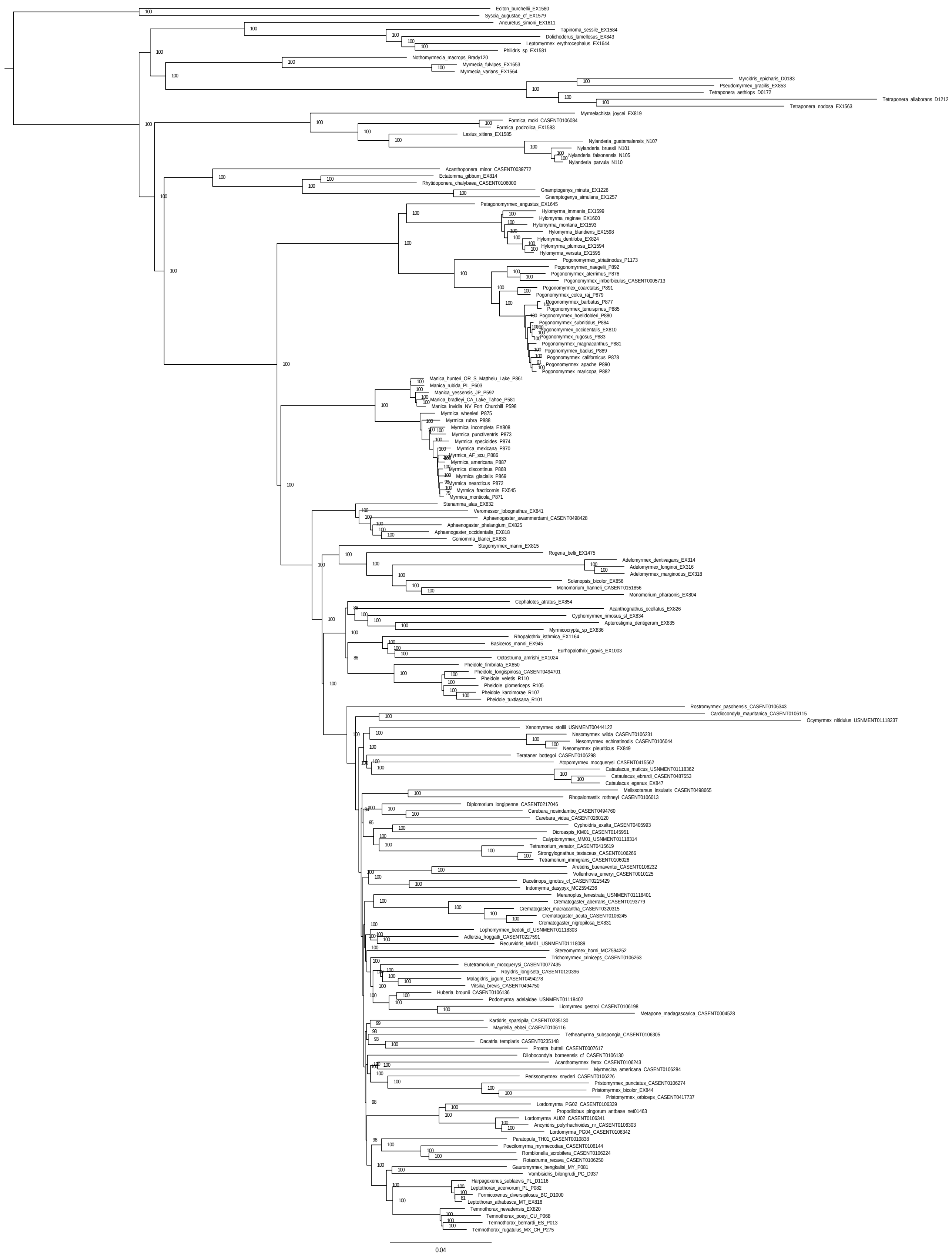


**Figure S14.** Results of maximum likelihood analysis of the ‘Myrmicinae UCEs’ dataset with IQTREE2, constrained to the Romiguier et al. 2022 topology. Statistical support at each node is in ultrafast bootstraps. Branch lengths are in units of substitutions per site.

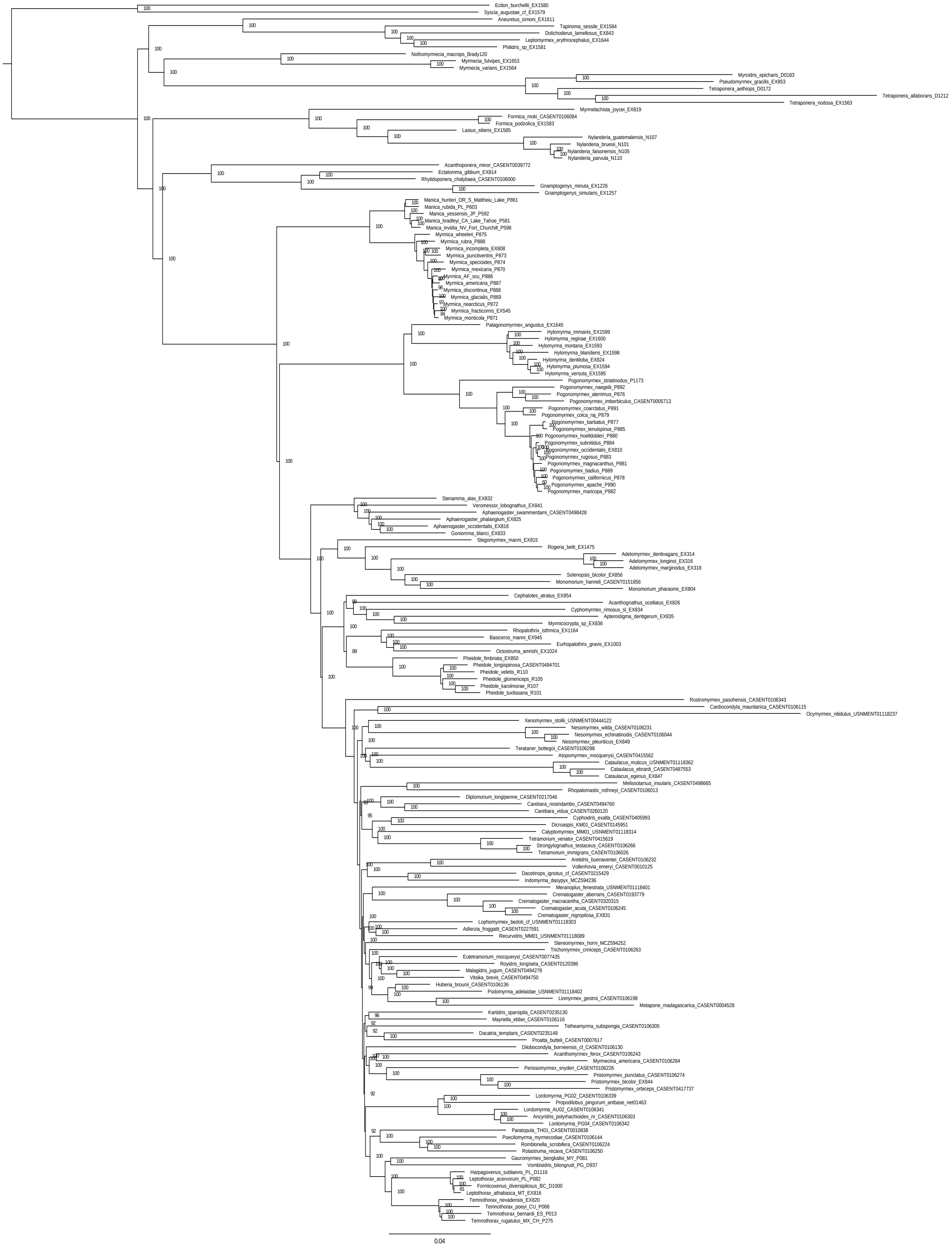


**Figure S15.** Results of maximum likelihood analysis of the 'Myrmicinae UCEs' dataset with IQTREE2, constrained to the Ward et al. 2015 topology. Statistical support at each node is in ultrafast bootstraps. Branch lengths are in units of substitutions per site.

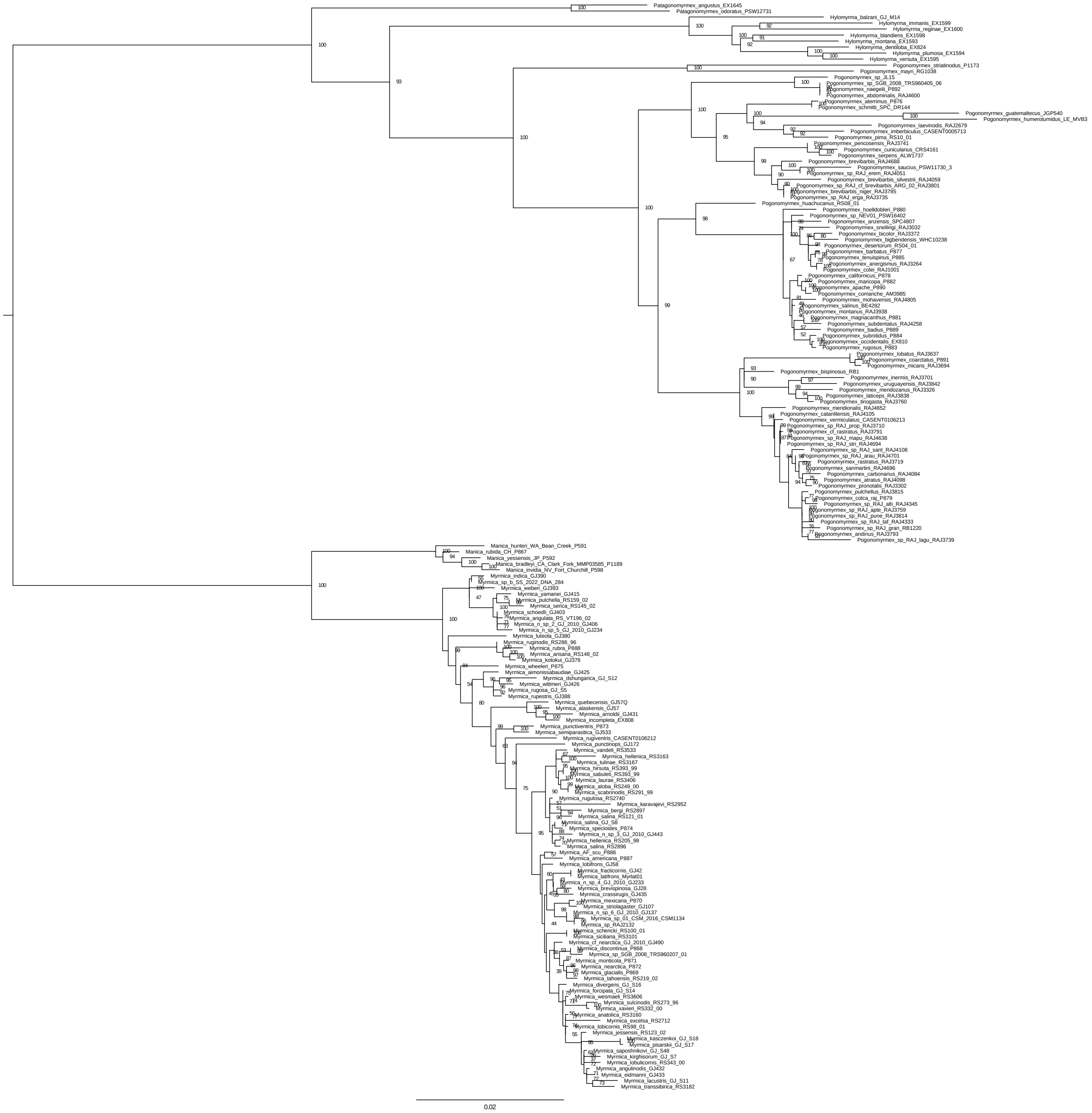


**Figure S16.** Results of maximum likelihood analysis of the 'Myrmicini Pogonomyrmecini' dataset with IQTREE2. Statistical support at each node is in ultrafast bootstraps. Branch lengths are in units of substitutions per site.

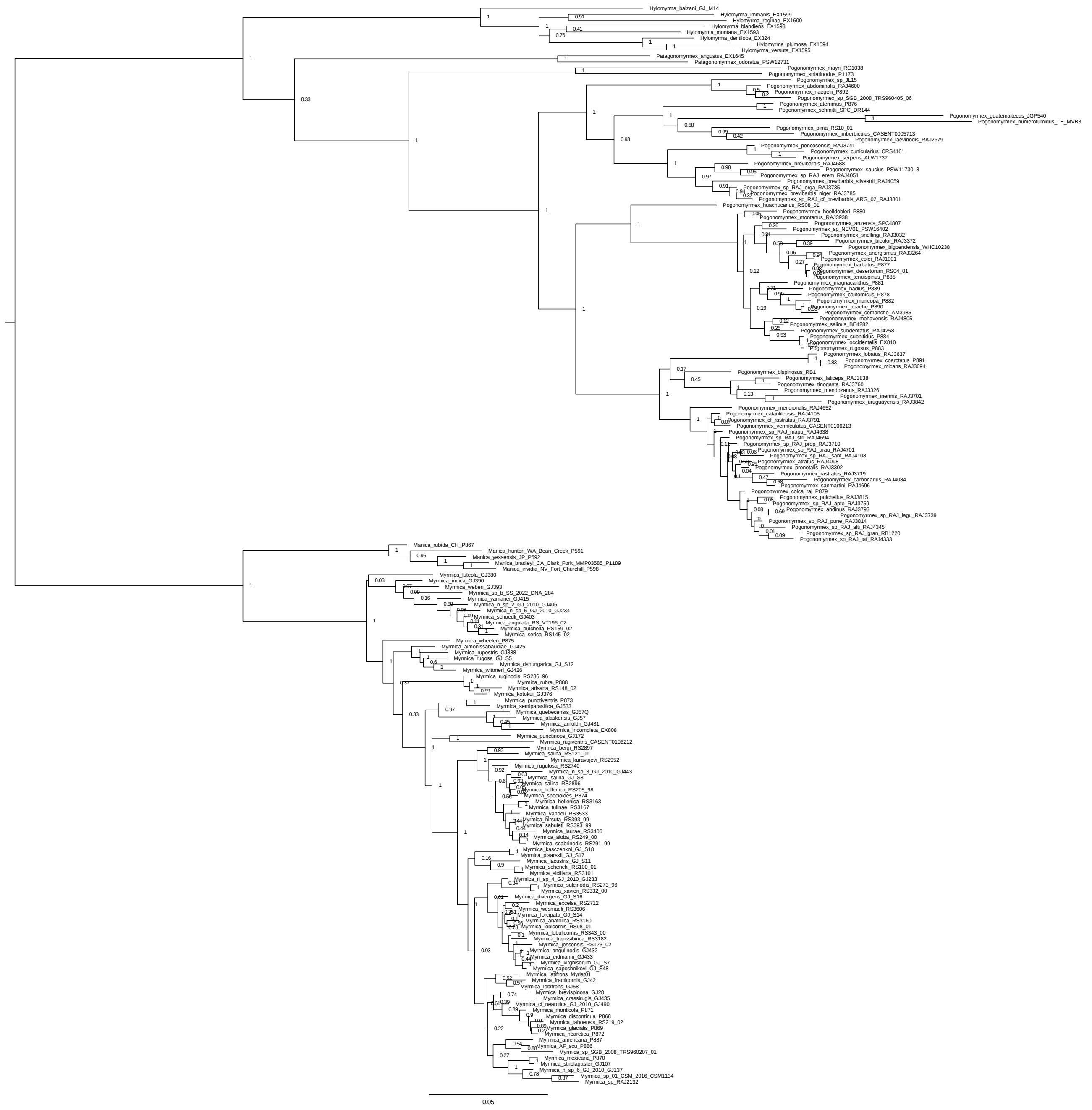

**Figure S17.** Results of Bayesian inference analysis of the ‘Myrmicini Pogonomyrmecini’ dataset with RevBayes. Statistical support at each node is in posterior probability. Branch lengths are in units of substitutions per site.

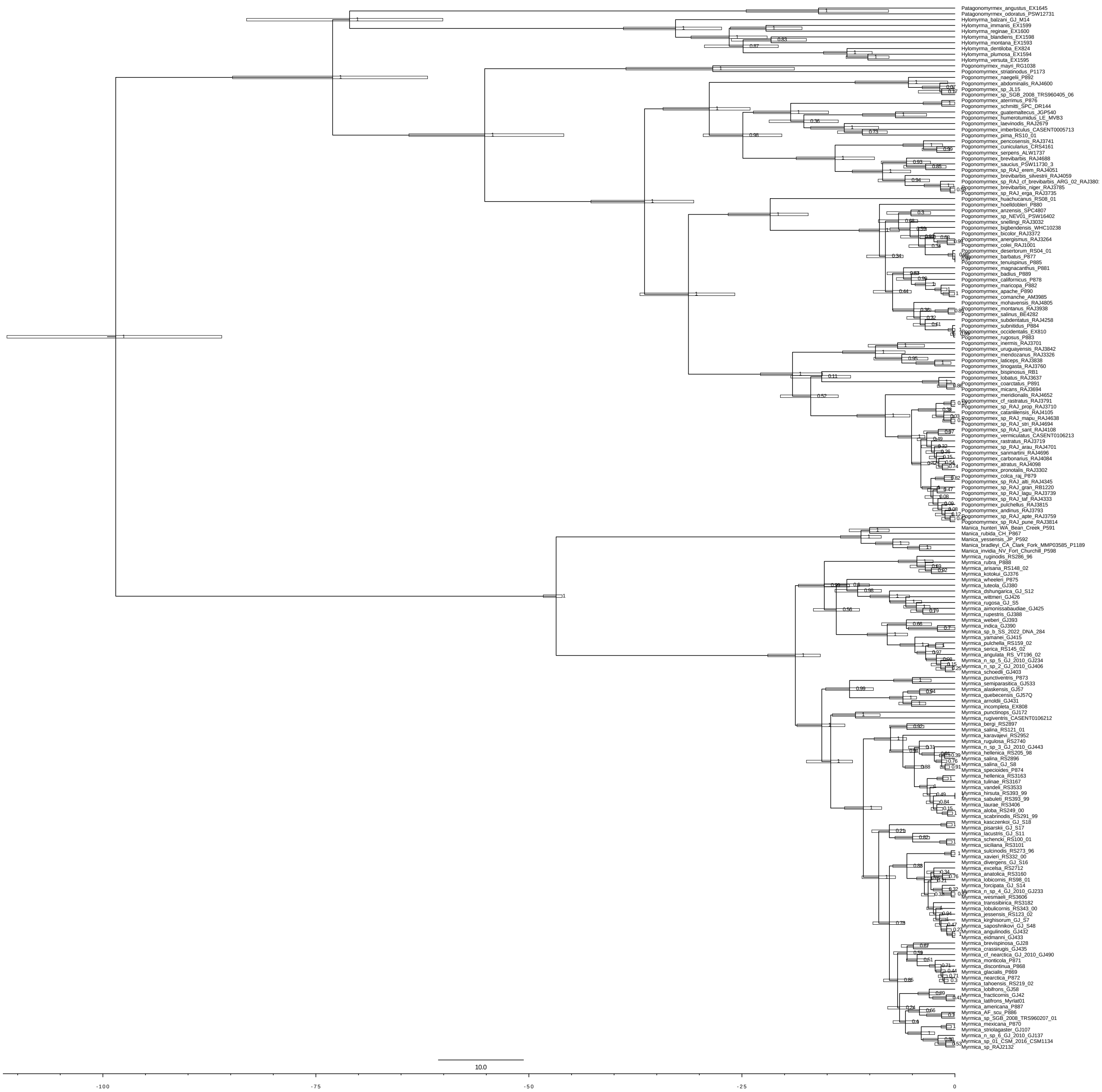

**Figure S18.** Results of divergence dating analysis of the ‘Myrmicini Pogonomyrmecini’ dataset with RevBayes. Bars around each node indicate 95% highest posterior density (HPD). Branch lengths are in units of one million years.

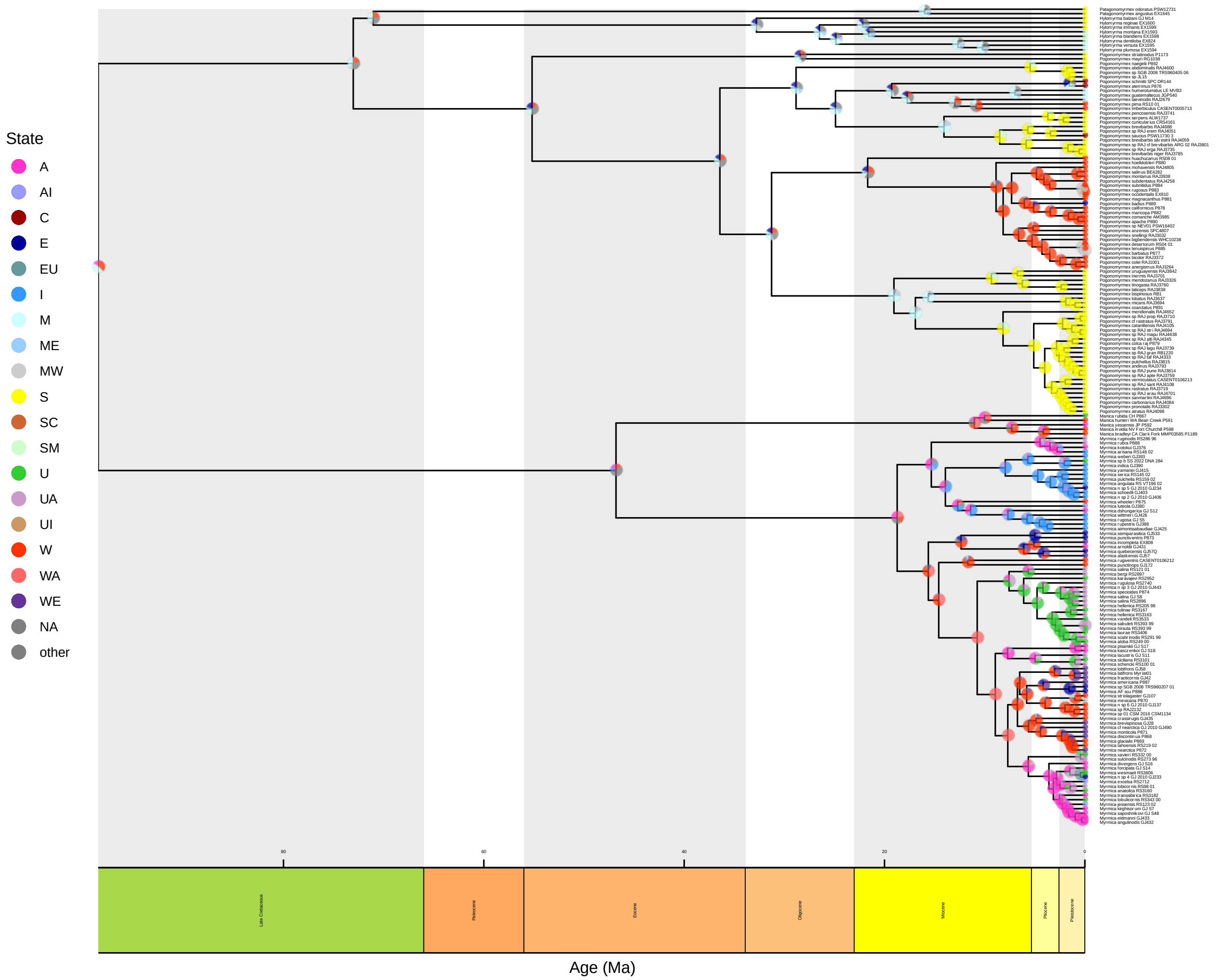


**Figure S19.** Results of biogeographical history ancestral state estimation analysis of the 'Myrmicini Pogonomyrmecini' dataset with RevBayes, without potential for overwater dispersal. Branch lengths are in units of one million years.

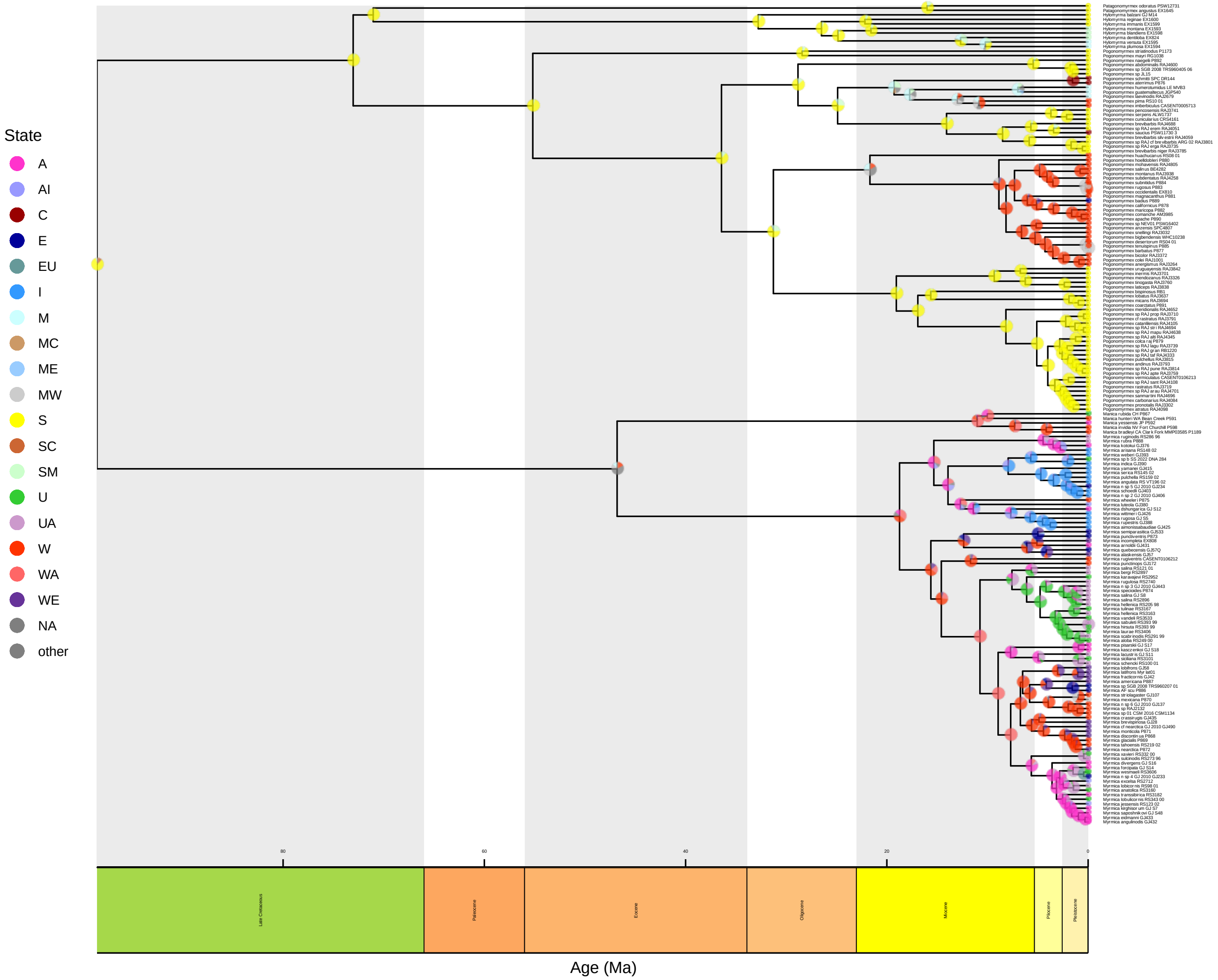


**Figure S20.** Results of biogeographical history ancestral state estimation analysis of the ‘Myrmicini Pogonomyrmecini’ dataset with RevBayes, with potential for overwater dispersal. Branch lengths are in units of one million years.
