## Supplementary file 3 for "Phylogenomics resolve the systematics and biogeography of the ant tribe Myrmicini and tribal relationships within the hyperdiverse ant subfamily Myrmicinae"

**Supplementary File S3.** Morphometric analysis of extant *Manica*, extant *Myrmica*, and the fossil species †*Manica andrannae*. Pairwise significance was calculated with a *t-test* at a significance level of $p < 0.05$.

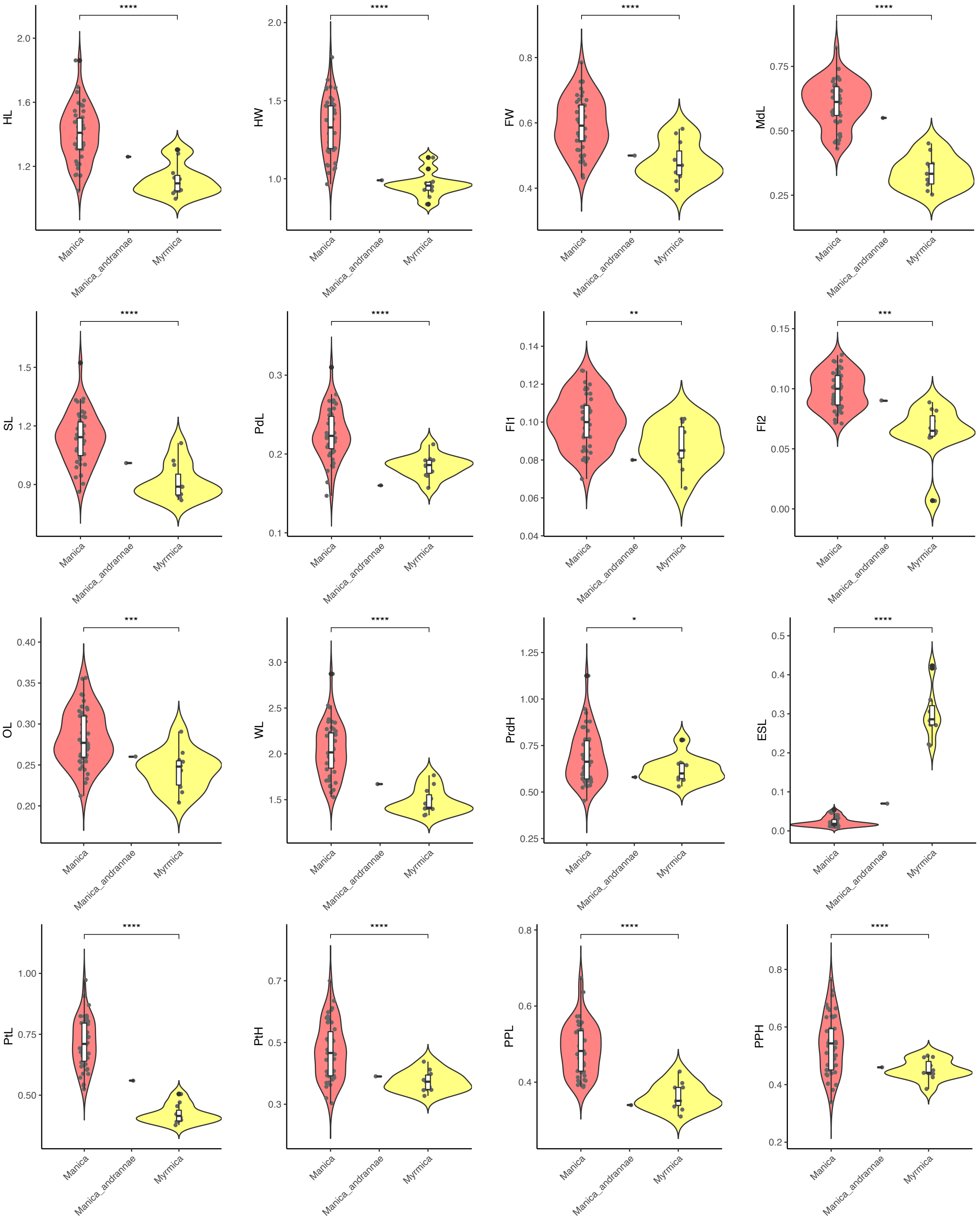

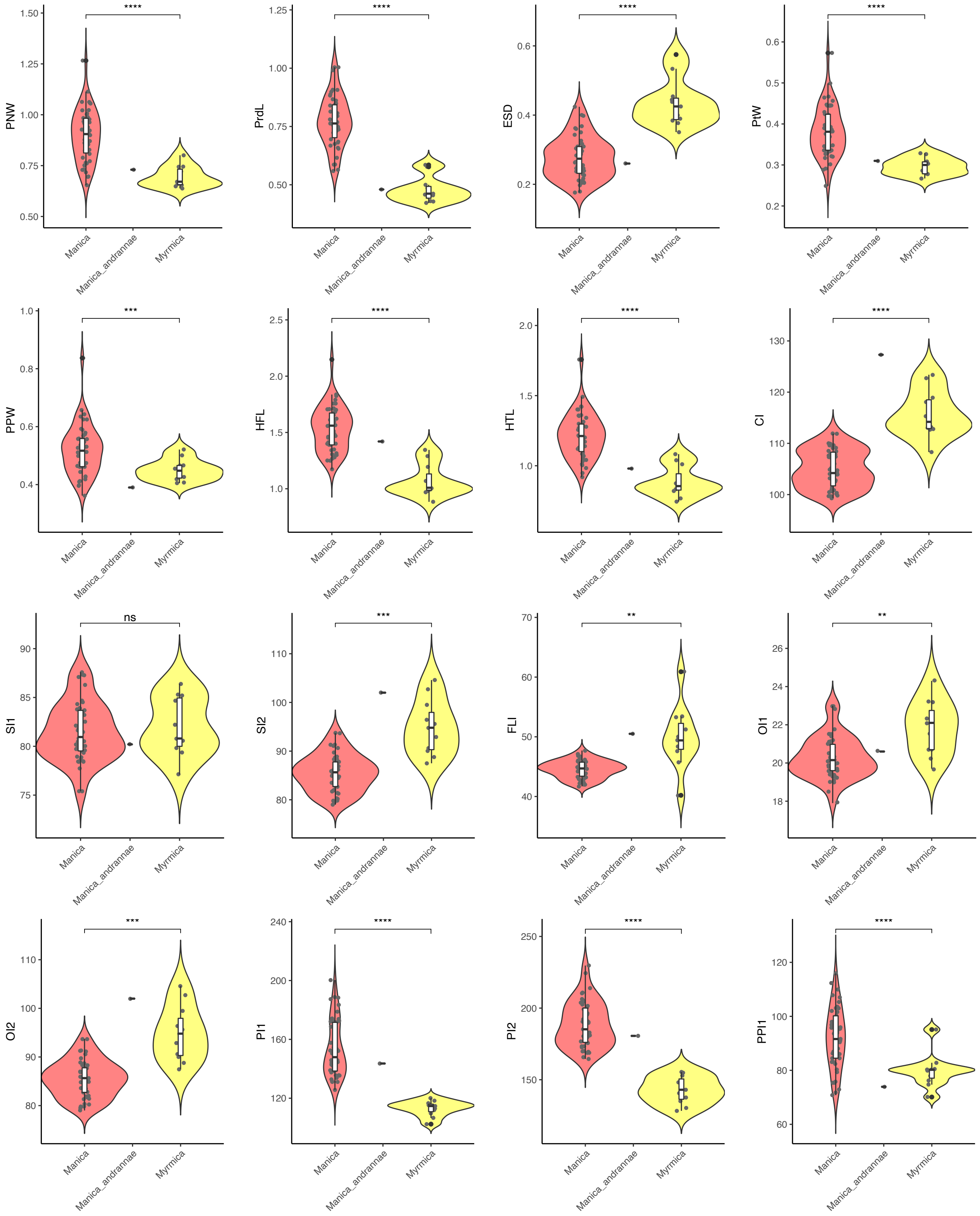
PNW
****
1.50
1.25
1.00
0.75
0.50
PrdL
****
1.25
1.00
0.75
0.50
ESD
****
0.6
0.4
0.2
PtW
****
0.6
0.5
0.4
0.3
0.2
PPW
***
1.0
0.8
0.6
0.4
HFL
****
2.5
2.0
1.5
1.0
HTL
****
2.0
1.5
1.0
CI
****
130
120
110
100
SI1
ns
90
85
80
75
SI2
***
110
100
90
80
FLI
**
70
60
50
40
OI1
**
26
24
22
20
18
OI2
***
110
100
90
80
PI1
****
240
200
160
120
PI2
****
250
200
150
PPI1
****
120
100
80
60
Manica
Manica_andrannae
Myrmica

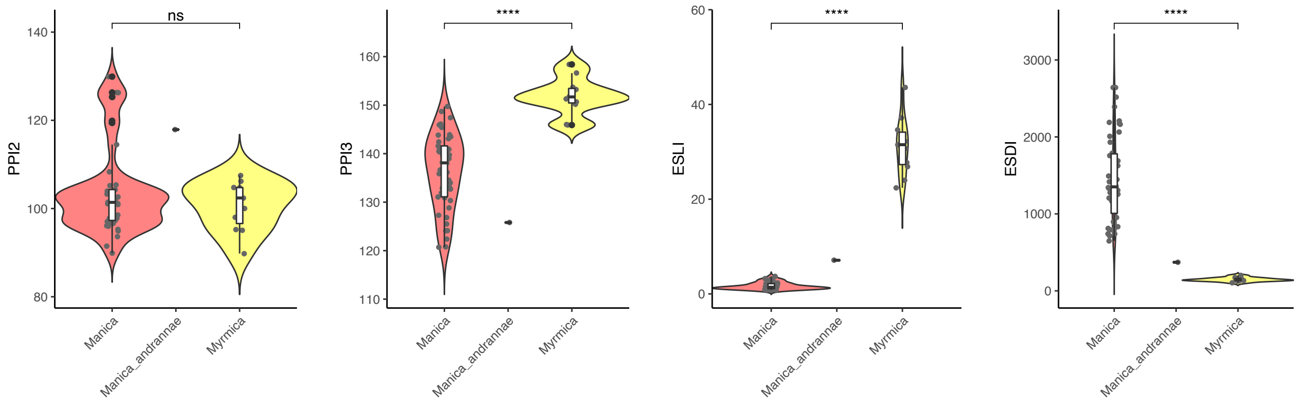

ns
****
****
****
PPI2
PPI3
ESLI
ESDI
Manica
Manica_andrannae
Myrmica
